# Model-Driven Elucidation of Lactose and Galactose Metabolism via Oxidoreductive Pathway in *Sungouiella intermedia* for Cell Factory Applications

**DOI:** 10.1101/2024.11.19.624258

**Authors:** Kameshwara. V. R. Peri, Ivan Domenzain, Hanna D Alalam, Abril Valverde Rascon, Jens Nielsen, Cecilia Geijer

**Affiliations:** Department of Life Sciences, Chalmers University of Technology, Gothenburg, Sweden; BioInnovation Institute, Ole Maaløes Vej 3, DK2200, Copenhagen, Denmark

**Author notes:** authors have contributed equally (Kameshwara. V. R. Peri, Ivan Domenzain).

**Keywords:** Non-conventional yeast, precision fermentation, Leloir pathway, galactitol, tagatose, flux balance analysis, genome-scale metabolic model

## Abstract

Converting industrial side streams into value-added chemicals using microbial cell factories is of increasing interest, as such processes offer solutions to reduce waste and production costs. However, developing new, efficient cell factories for precision fermentation remains challenging due to limited knowledge about their metabolic capabilities. Here, we investigate the lactose and galactose metabolism of the non-conventional yeast *Sungouiella intermedia* (formerly *Candida intermedia*), using knowledge-matching of high-quality genome-scale metabolic model (GEM) with extensive experimental analysis and determine its potential as a future cell factory on lactose-rich industrial side-streams. We show that this yeast possesses the conserved Leloir pathway as well as an oxidoreductive galactose catabolic route. Contextualization of RNAseq data into *Sint-GEM* highlights the regulatory mechanisms on the oxidoreductive pathway and how this pathway can enable adaptation to diverse environments. Model simulations, together with experimental data from continuous and batch bioreactors, indicate that *S. intermedia* uses upstream enzymes of the oxidoreductive pathway, in a condition-dependent manner, and produce the sugar alcohol galactitol as a carbon overflow metabolite, coupled to redox co-factor balancing during both lactose and galactose growth. Furthermore, the new metabolic insights facilitated the development of an improved bioprocess design, where an engineered *S. intermedia* strain could achieve galactitol yields of >90% of the theoretical maximum at improved production rates using the industrial side-stream cheese whey permeate as feedstock. Additional strain engineering resulted in galactitol-to-tagatose conversion, proving the versatility of the future production host. Overall, this work sheds new light on the intrinsic interplay between parallel metabolic pathways that shape the lactose and galactose catabolism in *S. intermedia*. It also demonstrates how a GEM combined with experimental analysis can work in synergy to fast-forward metabolic characterization and development of new, non-conventional yeast cell factories.

**Highlights:**

- An oxidoreductive pathway functions in concert with the Leloir pathway for galactose catabolism.
- GEM predicts that galactitol secretion enables efficient carbon overflow metabolism and maintains redox balance.
- Knowledge-matching of GEM with experimental results highlights cell factory potential.
- High galactitol yields and proof-of-concept tagatose production using whey permeate as feedstock.

## Introduction

Global challenges such as climate change, supply chain instability and loss of biodiversity pose threats to current production systems and result in failure to deliver and rising cost of chemical products. These challenges call for a stop of unsustainable production practices and a transition towards more innovative, sustainable and economically viable processes. Microbial fermentation offers flexible and robust production systems that are not restricted to specific geographical locations, climates and seasons, and active research and development have brought recent technological advancements to the field [1–3]. Microorganisms such as bacteria, yeasts and fungi can be used as “cell factories” for production of various metabolites of substantial societal and monetary value, including both bulk and speciality compounds. In addition to the well-known baker’s yeast *Saccharomyces cerevisia*e, widely used for production of various pharmaceuticals [4–7] and platform chemicals [8–11], non-conventional yeasts have also found applications due to their unique and desirable properties, e.g. *Yarrowia lipolytica* for lipids [4], *Pichia pastoris* for recombinant proteins [5, 6] and *Kluyveromyces lactis* for lactase production [7]. However, there is potential to explore biotechnologically useful functionalities in many more non-conventional yeasts, thereby expanding the range of chemicals that can be produced efficiently through precision fermentation.

The non-conventional yeast *Sungouiella intermedia,* formerly known as *Candida intermedia* [8], belongs to the CUG-Ser1 clade of ascomycetous yeasts and has the natural ability to utilize a wide range of different carbon sources [9]. Much of the initial work has focused on *S. intermedia*’s xylose assimilation and dwelled into its applicability as a potential cell factory for conversion of lignocellulosic hydrolysates into products [10–16]. Recent development of genome editing tools for this yeast, both marker-dependent [17] and marker-free editing using clustered regularly interspaced short palindromic repeats and CRISPR-associated protein 9 (CRISPR-Cas9) (manuscript in preparation), facilitate genotype-phenotype characterization and metabolic engineering. Using these genetic tools, we have also investigated its lactose and galactose metabolism, revealing novel regulatory, transcriptional and catabolic mechanisms involving three distinct gene clusters: the *LAC* cluster (*LAC12* lactose permease and *LAC4 β*-galactosidase), the *GAL* cluster (*GAL1*, *GAL10* and *GAL7* for galactose catabolism via the Leloir pathway), and a unique *GALLAC* cluster (second copies of *GAL1_2* and *GAL10_2*, transcriptional regulator *LAC9_2* and an aldose reductase *XYL1_2*) [18] (summarized in Fig. 1). Phenotypic analysis of gene deletion mutants revealed that the *GALLAC* cluster is indispensable for growth on lactose or galactose and signified the roles of Lac9 and Gal1_2 as regulators. Overall, this work revealed a complex interdependence between the gene products from the three clusters, where many of the proteins seem to exert both regulatory and enzymatic functions.

**Figure 1.**
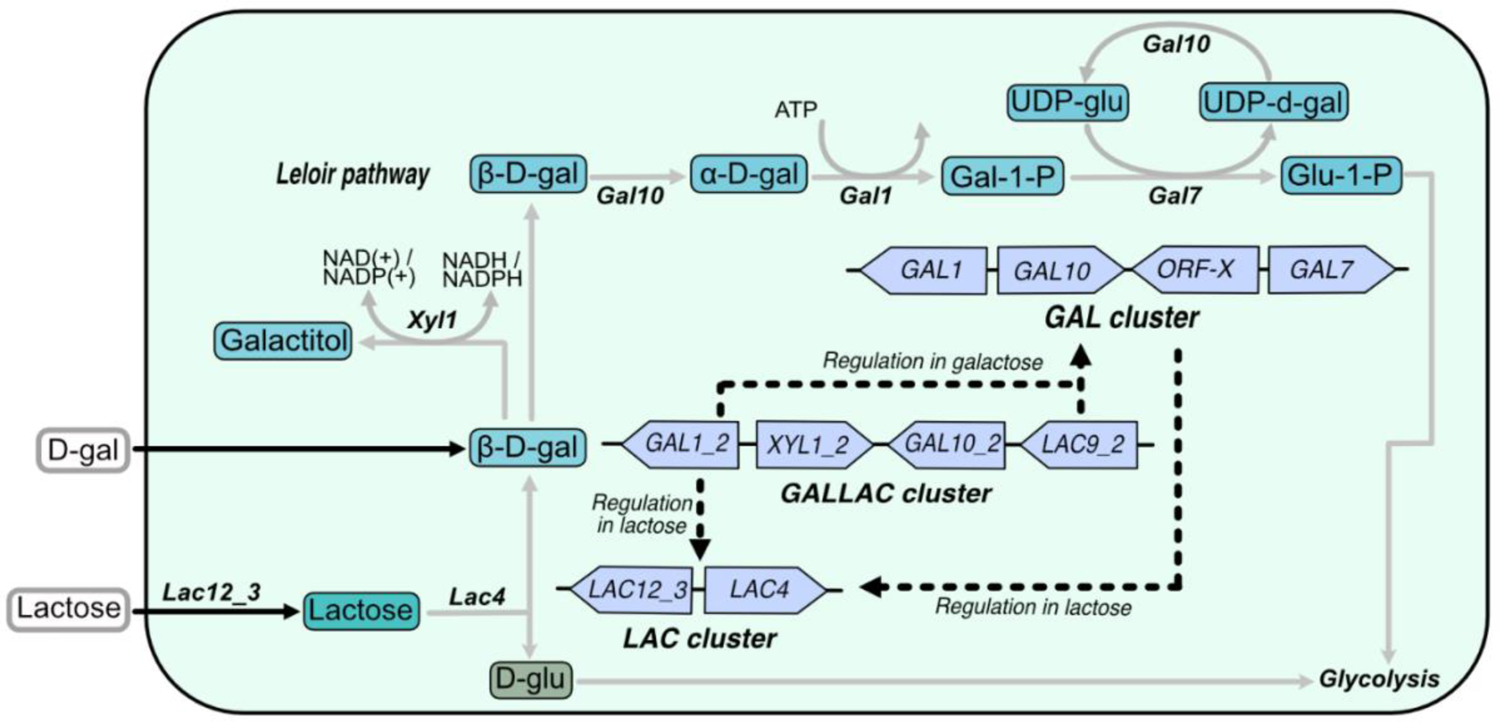
Depiction of the role of gene clusters in lactose and galactose metabolism in *S. intermedia*, adapted from [18]. Lactose metabolism starts with the uptake of lactose and subsequent hydrolysis to β-D-galactose. The *GALLAC* cluster gene *GAL1_2* exerts transcriptional regulation on *LAC* cluster genes (encoding uptake and hydrolysis of lactose) on lactose. *GAL1_2* along with *LAC9_2*, directly or indirectly, regulates *GAL* cluster (Leloir pathway enzymes encoding genes) expression on galactose. The *GAL* cluster has a regulatory control on the *LAC* cluster on lactose. Black solid rectangle depicts cell membrane. Transport is depicted by black solid arrows with the name of the associated transporter protein above it. Regulation is depicted by black dotted lines. Enzymatic conversion is shown by grey arrows and associated protein above or below. Abbreviations of metabolites are as follows – β-D-galactose (D-gal or β-D-gal); **α**-D-Galactose (**α**-D-gal); Galactose-1-phosphate (Gal-1-P); Glucose-1-Phosphate (Glu-1-P); UDP-glucose (UDP-gluc); UDP-galactose (UDP-gal) and D-glucose (D-glu).

Interestingly, deletion of the *GAL* cluster in *S. intermedia* resulted in extracellular accumulation of galactitol [18], a sugar alcohol with applications in the food industry as a precursor of the natural sweetener tagatose and in the pharmaceutical industry for the production of the anti-tumor agent di-anhydrogalactitol [19, 20]. Current production of galactitol relies on the chemical hydrogenation of galactose to galactitol, a process plagued by high costs and inefficiencies [21]. *S. intermedia’s* capacity to convert lactose into galactitol presents an opportunity to develop a new fermentation-based production process using whey, the abundant and lactose-rich side stream from the dairy industry, as substrate.

To engineer a new production host, which can reach high titers, yields and productivity in the bioproduction process, requires extensive understanding of its overall metabolism [22]. This can be achieved through a combination of experimental characterization and a systems level approach using a genome-scale metabolic model (GEM), which is a structured knowledge base of a cell’s metabolism in the form of a collection of *i)* genes in the genome of the organism, *ii)* metabolites in the cell and *iii)* reactions that they partake in [23]. Here, laws of thermodynamics and biochemical stoichiometry define and constrain the relations between such components, and thus, the cell itself [24, 25]. Furthermore, GEMs can be used as tools for quantitative simulation of metabolism under different genetic and environmental perturbations, enabling systematic exploration of the connection between genotypes and phenotypes [26, 27]. They can also facilitate a detailed understanding of a cells’ ability to convert substrates into products [28, 29]. For these reasons, GEMs are being extensively used to drive metabolic engineering projects in different application areas such as food [30], pharma [31–33] and human health [34–36].

With the overarching aim to understand the complex lactose and galactose metabolic network in *S. intermedia,* we curated and standardized a pre-existing model to generate a high-quality GEM, referred to as *Sint-GEM*. Combined with experimental results, we created a platform for probing the organism’s growth capabilities on lactose and galactose. Furthermore, we explored the potential of *S. intermedia* as a future cell factory for production of galactitol and its derivative tagatose; two metabolites with interesting characteristics for the food and pharma industries.

## Results and discussion

### *Sungouiella intermedia* can catabolize and grow on galactitol

In our previous work, we observed that deletion of the conserved *GAL* cluster in *S. intermedia*, which disrupts the Leloir pathway for galactose metabolism, results in slow growth on lactose along with production and extracellular accumulation of galactitol [18]. When we repeated the growth experiment and extended the cultivation time to 170 hours, we observed that the *galΔ* strain was not only capable of producing galactitol, but it could also consume it (Fig. 2a). Recent work from Rokas and colleagues shows that the *S. intermedia* type strain CBS 572 is among the 96 (out of 830) yeast or yeast-like strains that can grow on galactitol [37]. To test if this is a general trait for *S. intermedia,* we grew three *S. intermedia* strains (type strain CBS 572, our lab strain CBS 141442 and PYCC 4715) in minimal media with galactitol as the sole carbon source. Indeed, all three strains showed growth on galactitol, although the strain CBS 572 grew faster and reached higher OD levels than the other two strains within the time frame of the experiment (Fig. 2b). Nonetheless, these results collectively show that *S. intermedia* possesses a galactitol utilization pathway. This could also suggest that *S. intermedia* can catabolize galactose both through the canonical Leloir pathway and via galactitol as part of an alternative pathway, similar oxidoreductive pathways that have previously been reported in filamentous fungi *Aspergillus niger* and *Aspergillus nidulans* [38].

**Figure 2.**
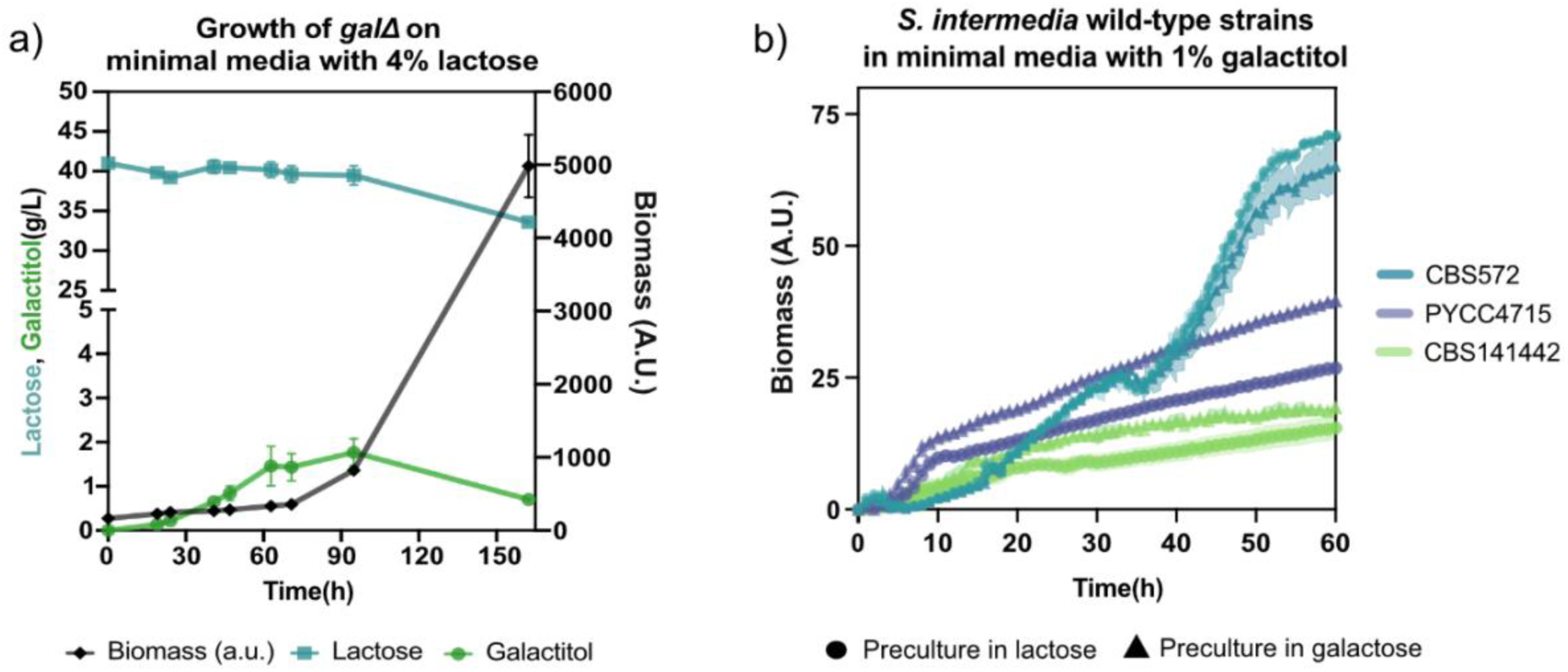
Galactitol production and consumption in *Sungouiella intermedia*. A) Growth curve, (biomass (a.u.) plotted in red with black squares plotted on right y-axis) and metabolites (lactose and galactose concentrations in g/L) plotted on left y-axis: lactose (in purple) and galactitol (in green) against Time (in hours on x-axis), of the *galΔ* mutant grown in minimal media containing 4% lactose as carbon source. Data is represented for biological duplicates with error depicted by shaded region for biomass and error bars for metabolites. B) Growth curves of three *S. intermedia* strains (CBS 572 (dark green); PYCC 4715 (purple); CBS 141442 (light green)) on minimal media with 1% galactitol as carbon source. Strains were precultured either in lactose (filled circle) or galactose (filled triangle) before starting the main culture in Growth Profiler960 (Enzyscreen, The Netherlands). These carbon sources were chosen to avoid putative glucose repression of the galactitol-catabolic genes prior to cultivation. Biomass is represented in green values (A.U.) on Y-axis against time in hours on X-axis. Data is represented for biological triplicates with error depicted by shaded region for biomass.

### *Sint-GEM* predicts metabolites belonging to an oxidoreductive pathway

To investigate the presence of an oxidoreductive pathway in *S. intermedia*, and to elucidate the potential benefits of having two parallel catabolic pathways for galactose catabolism, we developed a GEM for *S. intermedia*, hereafter referred to as *Sint-GEM*.

An initial draft model of *S. intermedia’* metabolism was generated in a previous study [39], following an automated process, based on the consensus GEM for *S. cerevisiae* Yeast8 [40] and orthology searches across 332 sequenced yeasts and 11 other fungi (Fig. 3a). Assessment of the representation of lactose metabolism in the initial GEM revealed a lack of the lactose uptake step, which can be attributed to the inherited use of Yeast8 as a template network (*S. cerevisiae* lacks the gene for lactose uptake). This reaction step, for which we have previous experimental evidence [18], were therefore introduced into the GEM (Fig. 3b).

**Figure 3.**
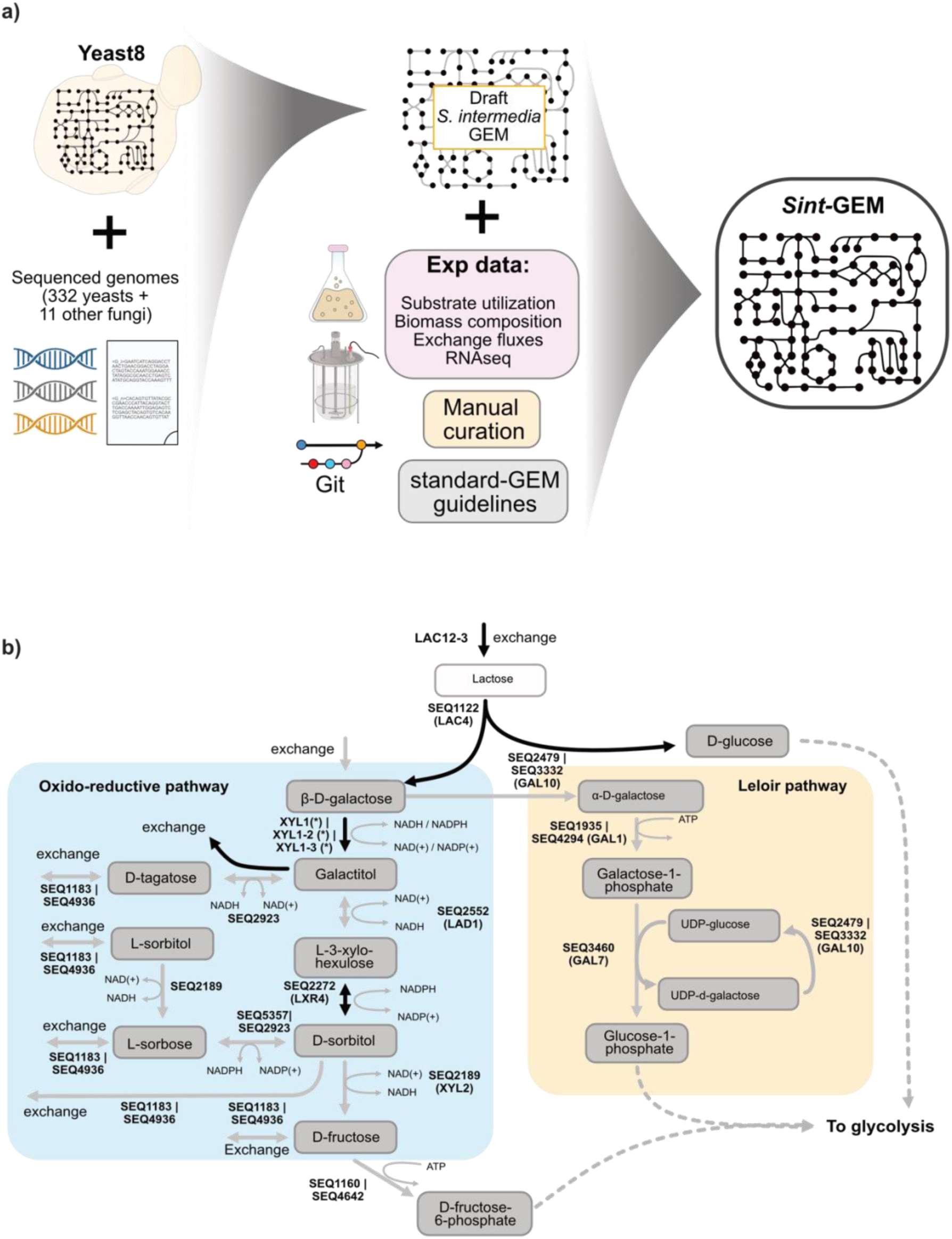
Construction of a GEM for *S. intermedia*. A) Reconstruction of the *Sint-GEM* using draft model for *S. intermedia* based on yeast8 and gene orthology predictions. Data-driven manual curation and git-based version control yielded a high-quality GEM adhering to standard modeling practices. *Sint-GEM* consists of 1070 genes, 3991 reactions, 2759 metabolites and 14 cellular compartments. B) Representation of lactose and galactose metabolism in *Sint-GEM*. Curation of the oxidoreductive pathway in *Sint-GEM* involved addition of L-xylo-3-hexulose reductase reaction connecting L-xylo-3-hexulose with D-sorbitol (indicated with black arrows). Additionally, cofactor usage of galactose reductase was incorporated into the model according to experimental results.

The initial model draft included all metabolites and enzymes associated with the conserved Leloir pathway, as well as the galactose-to-galactitol conversion step catalyzed by an aldose reductase. *S. intermedia* possesses no less than three paralogous genes encoding aldose reductases, previously identified as xylose reductases [15]. Among these, Xyl1_2, is encoded from the *GALLAC* cluster that is essential for growth on lactose and galactose [18]. All three enzymes and their cofactor specificities were incorporated in the model, where Xyl1_2 displays dual redox (NADH/NADPH) cofactor specificity (Fig. S1), while Xyl1_1 and Xyl1_3 are strictly NADPH dependent [15]. As extracellular galactitol accumulation and subsequent consumption were observed in the *galΔ* mutant strain (Fig. 2a), an exchange reaction for galactitol transport/diffusion into and out of the cell was also integrated into the model.

Notably, the model already included the reaction steps for converting galactitol to L-xylo-3-hexulose, catalyzed by *LAD1*-encoded galactitol dehydrogenase, as well as the conversion of D-sorbitol to D-fructose by the *XYL2*-encoded D-sorbitol dehydrogenase. To complete the oxidoreductive pathway, the reaction converting L-xylo-3-hexulose to D-sorbitol by L-xylo-3-hexulose reductase was added. Here, the *LXR4* gene product was assigned this enzymatic activity, based on ortholog predictions between *S. intermedia* and filamentous fungi shown to display an oxidoreductive pathway for galactose catabolism [41]. Together, these reaction steps connect all metabolites from β-D-galactose to D-sorbitol, and further into D-fructose and D-fructose-6-phosphate for glycolytic assimilation (Fig. 3b). In addition to the steps mentioned above, *Sint-GEM* contained additional reactions involving possible oxidoreductive pathway intermediates, including the conversion of galactitol to tagatose by sorbitol dehydrogenase encoding *SOU2* (Seq_2923) and L-sorbitol to L-sorbose by *YALI0B16192G* (Seq_5357) or *SOU2*.

To further refine the predictive capabilities of *Sint-GEM*, data on biomass composition and consumption/secretion rates of metabolites from continuous cultivations were incorporated and utilized to fit the model’s energy requirements. *Sint-GEM* development is tracked following standard-GEM guidelines [42] and can be found at: https://github.com/SysBioChalmers/cint-GEM.

### Transcriptional insights support the *Sint-GEM*-predicted oxidoreductive pathway

Attempting to elucidate the network topology of the oxidoreductive pathway, we started by contextualizing the differential gene expression observed during lactose or galactose growth compared to glucose growth (reported previously in [15]), in *Sint-GEM*. An overview of the transcriptional profiles is shown in Figure 4. Metabolic contextualization of transcriptional data depicted a cellular condition in which expression of the aldose reductase isoforms Xyl1_2 and Xyl1_3 pulls flux toward conversion of galactose into galactitol. Galactitol can be secreted out of the cell, as observed in the *galΔ* strain, or further catabolized by the expressed enzymes Lad1 and Xyl2, to provide additional glycolytic flux. Additionally, strong downregulation of genes encoding for transporters of other oxidoreductive metabolites (D-tagatose, L-sorbose and D-sorbitol) can minimize the loss of additional carbon flux. Overall, based on gene expression profiles, flux through the oxidoreductive pathway in *Sint-GEM* suggests a possible coordinated effort in maximizing carbon assimilation.

**Figure 4.**
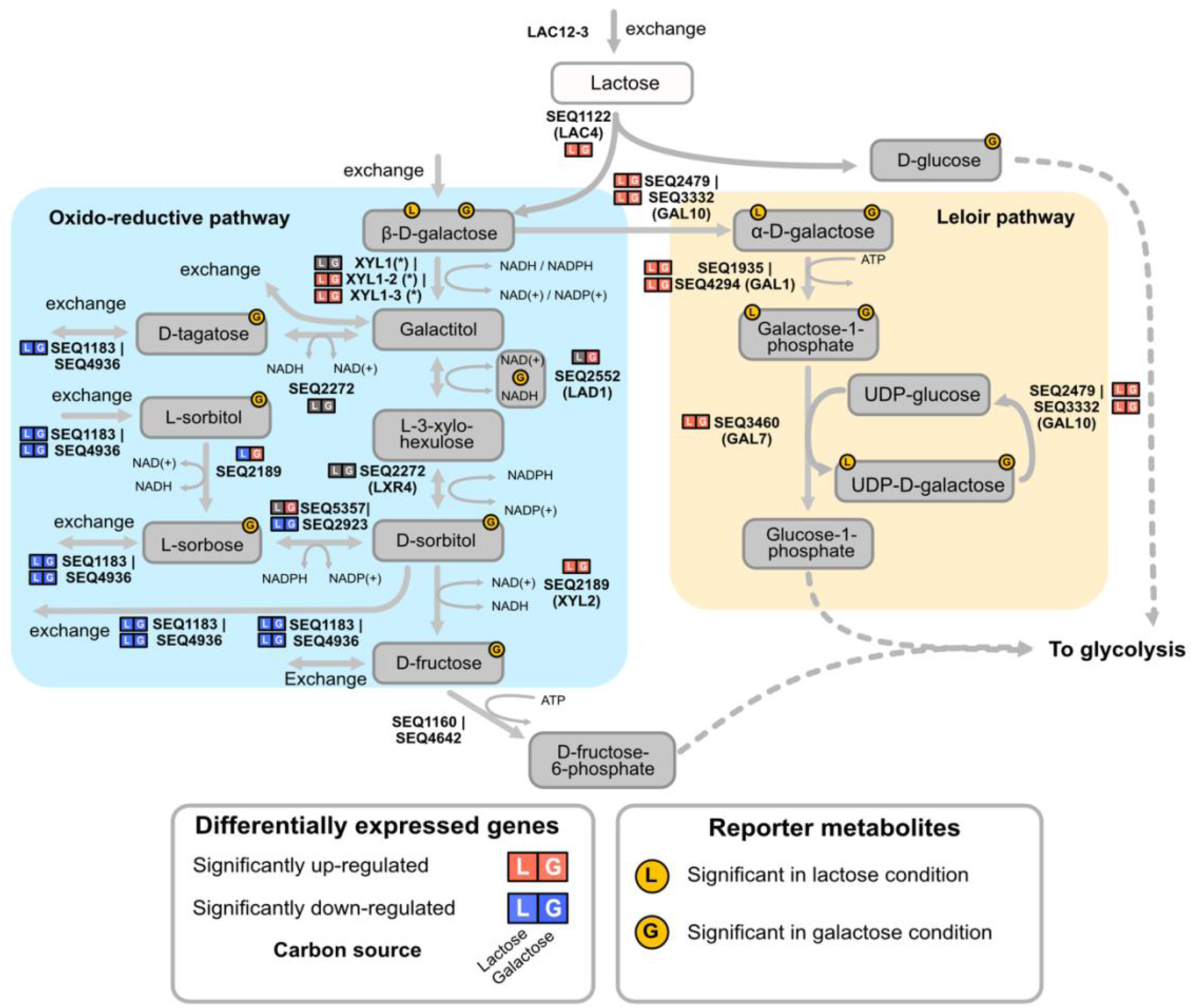
Transcriptional insights support activity of the *Sint-GEM*-predicted oxidoreductive pathway during growth on lactose and galactose. Differential expression profiles and reporter metabolite analysis of genes and pathway intermediates of the Leloir pathway and the hypothesized oxidoreductive pathway in S. intermedia wild-type strain on galactose and lactose. Differential expression of genes is depicted by squares indicating directionality (red = upregulation; blue = downregulation) and the utilized carbon source (indicated by L = lactose and G = galactose). Similarly, reporter metabolites are represented by yellow circles indicating the carbon source in which the metabolite is being characterized.

Additionally, we performed reporter metabolite analysis [43], again utilizing the results from the differential gene expression analysis on growth on lactose or galactose compared to glucose [15]. Results for both carbon sources showed that 4 out of the 6 metabolites participating in the Leloir pathway become significant nodes of transcriptional regulation (adjusted p-value <0.001), i.e. indicating strong and coordinated transcriptional up/down-regulation of genes associated with these metabolites (Fig. 4). In contrast, only β-d-galactose, the first metabolite entering the oxidoreductive pathway, was found among the top reporter metabolite on both carbon sources. Several other metabolites in the oxidoreductive pathway (NAD(+/H), D-sorbitol, L-sorbose, L-sorbitol and D-tagatose) were found to be nodes of significant regulation only on galactose. These results indicate that lactose availability seems to favor activation of the Leloir pathway to metabolize the galactose moiety, while the glucose moiety represents a direct contribution to glycolytic flux in the cell. In contrast, activation of both the Leloir and oxidoreductive pathways is coupled to catabolism of galactose as sole carbon source for both energy and redox balance purposes.

### Flux balance analysis suggests conditional use of the oxidoreductive pathway

Next, using flux balance analysis (FBA) [44], we verified the connectivity and functionality of lactose and galactose metabolism in *Sint-GEM* by predicting cellular growth capabilities on each carbon source. Simulated flux distributions in both carbon sources suggest that the Leloir pathway is the stoichiometrically optimal route for galactose catabolism [18]. However, when using galactose as carbon source and forcing assimilation via the oxidoreductive pathway (by blocking conversion of α-D-galactose into galactose-1-phosphate), only a slight decrease in the simulated biomass yield (3.14%) was observed, a phenotype mainly explained by 2.27-fold increase in the divergence of glycolytic flux toward the pentose-phosphate pathway (PPP) for NADPH regeneration, in comparison to the optimal Leloir utilizing phenotype (Suppl. Table 1). Thus, FBA results indicate that, although the Leloir pathway is the preferred route for galactose metabolism, the oxidoreductive pathway can also carry flux and stoichiometrically support cellular growth on both carbon sources.

To further explore the conditions in which *S. intermedia* carries flux through the oxidoreductive pathway, we probed the metabolic landscape of the cell, following a random flux sampling procedure [45]. Both galactose and lactose were used as carbon source under three different scenarios of cellular growth: carbon-, oxygen-, and nitrogen-limited conditions, and a total of 10,000 FBA simulations with randomized objective functions were sampled for every condition. Each of these flux distributions simulated a different attainable phenotype for the cell at the specified constraints, as illustrated by Figure 5a.

**Figure 5.**
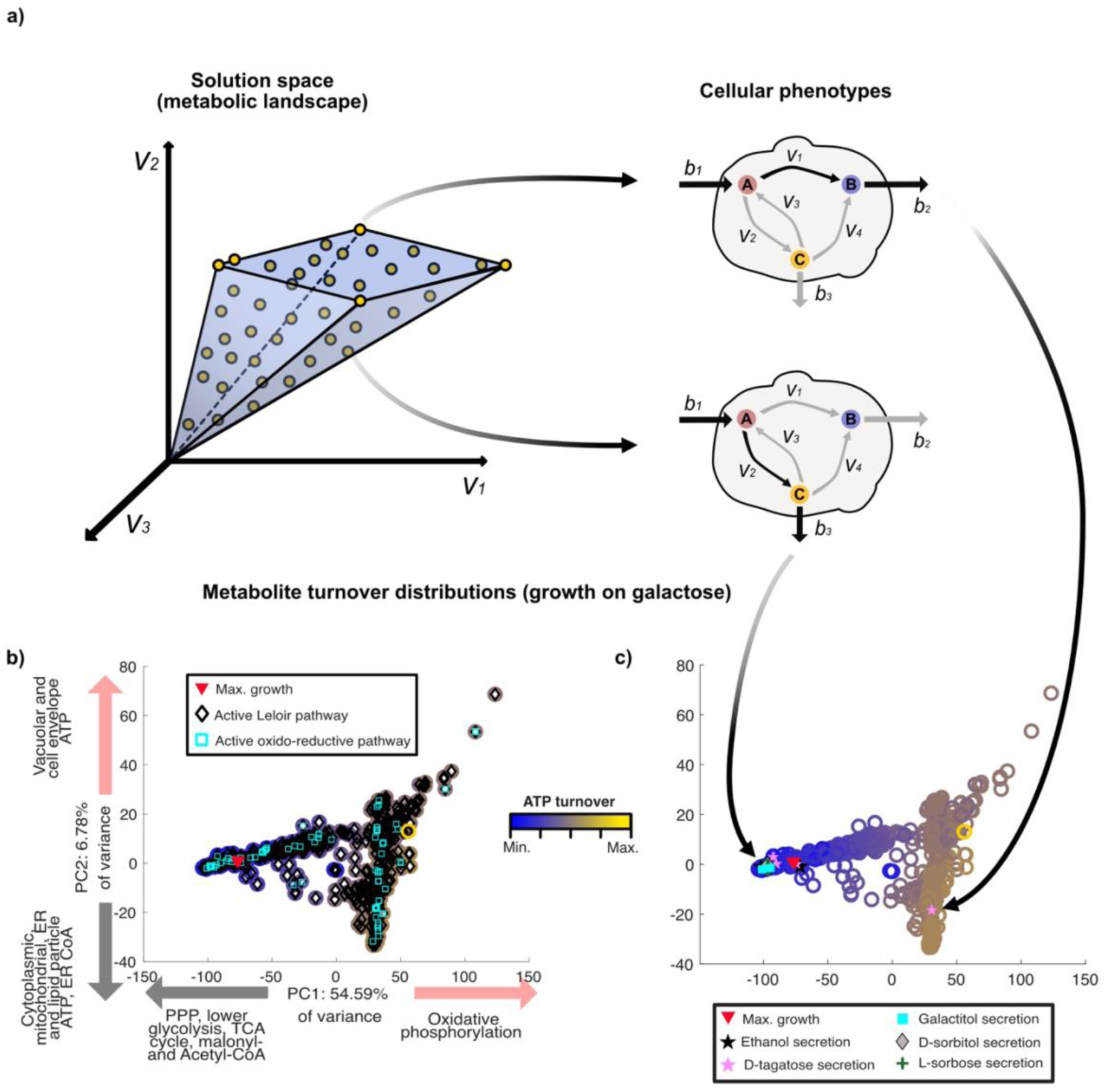
Probing the solution space of *Sint-GEM* enables exploration of the metabolic flux simulations of *S. intermedia* cells. A) FBA simulations with random objective function are computed, allowing the exploration of attainable cellular phenotypes given a set of specified constraints. (b-c) Principal component analysis of flux distributions. Metabolite turnover distributions are computed from flux distributions and dimensionally reduced using principal component analysis. Sampled FBA simulations are mapped into PC1-PC2 space. Arrows and legend next to PCs axis represent the metabolites (also related to pathway or subcellular localization) whose turnover rates are the top contributors to the respective PC in both positive and negative directions. Both (b) and (c) represent the same 10,000 simulations of cellular growth under carbon-limited conditions with galactose as sole carbon source). Color of each sample (rings) represents the normalized total intracellular ATP turnover (dark blue for minimal values, bright yellow for maximal ones). B) The use of the Leloir pathway and/or the oxidoreductive pathway for galactose catabolism is mapped onto each of the sampled distributions. Simulations achieving the maximum optimal growth are marked by a red inverted triangle. C) FBA simulations secreting ethanol, D-tagatose or galactitol are mapped into PC1-PC2 space of the sampled metabolite turnover distributions.

Flux distributions were transformed into metabolite turnover distributions followed by a dimensionality reduction using principal component analysis (PCA) (see materials and methods for further details), to obtain a global view of the *Sint-GEM* solution space. Figure 5b illustrates that, under carbon-limited conditions with galactose as the carbon source, the spread of flux distributions through the Leloir pathway and the oxidoreductive pathway over the *Sint-GEM* solution space are overlapping. In fact, this was observed across all limiting conditions explored by random sampling, indicating that either use of the pathways is not associated to a particular metabolic profile or a specific compartment in the cell (Fig. S2a-f). As expected, according to the computed optimal FBA simulations under both carbon sources (Suppl. Table 1), the Leloir pathway carries flux for assimilation of galactose in most simulations across all conditions (>99% in all cases, see Suppl. Table 2). Further analysis revealed that dual galactose pathway usage was present in a high number of simulations, especially in the oxygen-limited and nitrogen-limited cases with galactose as carbon source, and the oxygen-limited case on lactose (100%, 42.64% and 60.9% of the 10,000 simulations explored for each case). Only a small fraction of the simulated flux distributions (<1%) demonstrate exclusive use of the oxidoreductive pathway (Suppl. Table 2). Interestingly, random sampling analysis across all conditions suggests that >95% of the flux distributions through the oxidoreductive pathway show a strong preference for NADH as cofactor for the aldose reductase Xyl1_2. Dual cofactor usage was predicted for ∼42% of the simulations, while NADPH was exclusively preferred in only a few cases across different conditions, with the exception of galactose-limited conditions where NADPH preference was observed in about 45% of cases.

### Aldose reductase Xyl1_2 ensures redox cofactor balance in *Sungouiella intermedia*

The dual cofactor specificity of the aldose reductase Xyl1_2 is a peculiar characteristic, as most other known aldose reductases are strictly NADPH dependent [46, 47]. We used *Sint-GEM* to understand better its potential role in lactose and galactose metabolism. To this end, we blocked the Leloir pathway, and the flux distribution for growth on galactose showed that NADH is the stoichiometrically optimal cofactor for galactitol formation. Next, we blocked both the Leloir pathway and NADH usage, to evaluate the potential impact of exclusive NADPH utilization on growth outcomes. This revealed a significant compromise of glycolytic flux for NADPH regeneration via the Pentose Phosphate Pathway (PPP). The 2.8-fold increase in PPP flux compared to the basal distribution in the wild-type growing on galactose suggested a compromise in overall biomass yield (Suppl. Table 3 and 4).

To explore the impact of NADPH use by the aldose reductase on a genome-wide scale, we analyzed the correlation between NADPH oxidation rate and metabolite turnover across all simulated conditions (carbon, oxygen, and nitrogen limitations, using galactose or lactose as carbon sources) (Suppl. Tables 3 and 4). In carbon- and oxygen-limited conditions using galactose as a carbon source, NADPH consumption in the aldose reductase step showed a significant positive correlation with turnover rates of TCA cycle intermediates. Additionally, we found a significant negative correlation between NADPH usage and turnover of Leloir pathway metabolites under carbon-limited conditions for both galactose and lactose. Under oxygen-limited conditions with galactose, NADPH oxidation in this step was linked to higher production of long-chain phospholipids in vacuolar and Golgi membranes. Similarly, in lactose-limited conditions, NADPH consumption was associated with increased turnover of very-long-chain ceramides in the ER and Golgi, connected to lipid biosynthesis and membrane composition [48]. Overall, these results emphasize the significance of Xyl1_2’s cofactor flexibility, as its ability to also use NADH helps maintain cellular balance and prevents potential disruptions to the cell’s macromolecular composition due to excessive NADPH oxidation.

### Model and quantitative fermentation unravel galactitol and ethanol secretion under nutrient-limited conditions

As shown experimentally, the *S. intermedia galΔ* strain accumulates galactitol extracellularly when grown on lactose (Fig. 2a) [18]. This motivated analysis of secretion fluxes across all simulations explored through random sampling, aiming to identify the conditions under which other metabolites in the oxidoreductive pathway might be secreted from the cell. Simulations of the wild-type strain secreting the metabolites galactitol, D-sorbitol, L-sorbose and D-tagatose were mapped onto the PCA plots of the randomly sampled metabolite turnover distributions (Fig. S3). We also mapped ethanol, as this is a metabolite that is formed during oxygen-limited conditions in many yeasts, including *S. intermedia* [15].

#### Carbon-limiting conditions

Under carbon-limiting conditions in galactose and lactose, a very minor fraction of the random sampling simulations displayed secretion of any of the metabolites of interest (less than 1% for all cases, Table 1). Similarly, lactose-limited chemostat cultivations of the wild-type strain showed no accumulation of the metabolites when tested at two different dilution rates (data not shown). This suggests that the strain grew under optimal conditions, exhibiting primarily respiratory metabolism.

**Table 1.** Percentage of simulations for each secreting ethanol and oxidoreductive pathway intermediates in FBA random sampling using *Sint-GEM* at different limiting conditions. 10,000 simulations were run for each combination of carbon source and nutrient limitation.

| Metabolite | Galactose |  |  | Lactose |  |  |
| --- | --- | --- | --- | --- | --- | --- |
|  | C-limited | O2-limited | N-limited | C-limited | O2-limited | N-limited |
| Ethanol | 0.13% | 94.84% | 0% | 0.24% | 100% | 0 |
| Galactitol | 0.8% | 60.51% | 0.33% | 0.11% | 100% | 42.93% |
| D-tagatose | 0.19% | 0% | 0% | 0.13% | 0% | 0 |
| D-sorbitol | 0.04% | 0% | 0% | 0.09% | 0% | 0 |
| L-sorbose | 0.03% | 0% | 0% | 0.13% | 0% | 0 |

FBA simulations for growth under galactose-limiting conditions predicted complete respiration of the carbon source with no secretion of byproducts (Fig. 6a). In contrast, growing the wild-type strain in batch reactors using galactose as a carbon source resulted in significant levels of galactitol production. Analysis of the media composition and gas inputs and outputs in the reactor revealed that the time interval of highest rate of galactitol production coincided with maximum cellular growth, the highest rate of galactose consumption and the highest (but non-depleting) oxygen uptake rates (between 36-60 hours – marked by dotted lines, Fig. 6b, c). During extended cultivation (60 hours, Fig. 6b), secreted galactitol began to be consumed as galactose levels neared depletion in the media. This switch from production to consumption suggests that cells may be reaching primary carbon-limiting conditions, aligning with the predictions shown in Fig. 6a and the random sampling results under galactose-limited conditions (Table 1).

**Figure 6.**
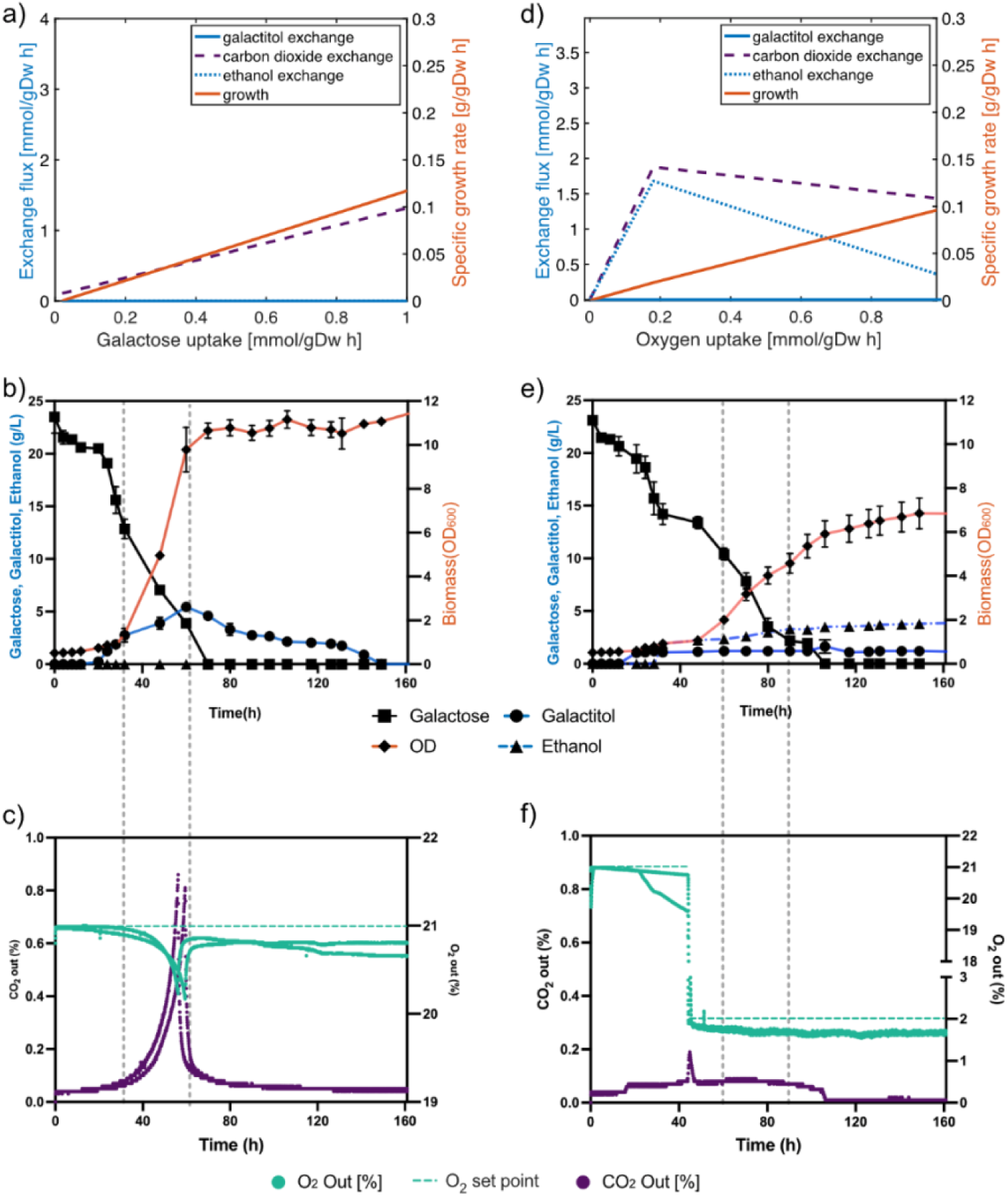
Model and quantitative fermentation unravel galactitol and ethanol secretion under nutrient-limited conditions. A) Predicted growth rate and metabolites exchange at varying galactose uptake rates in *Sint-GEM*. Exchange flux (in mmol/gDW h) of galactitol (blue solid), ethanol (dotted light blue), and CO_2_ (black dash line) shown on left y-axis, with specific growth rate (solid red line) on the right y-axis, are plotted against galactose uptake rate (in mmol/gDW h). B) Cell density and extracellular metabolite concentration over time for wild-type *S. intermedia* grown on batch reactors with standard aeration levels (21%). Biomass (OD, represented with diamonds), extracellular concentration of galactose (squares), galactitol (circles) and ethanol (triangles). C) Percentage of O_2_ (aqua) and CO_2_ (purple) measured at the air outlet along the cultivation time in batch reactors with standard aeration levels (21% O_2_ as set point, indicated as an aqua-colored dotted line). Vertical grey dotted lines delimit the time span with the highest measured growth and galactose consumption rates. D) Predicted growth rate and metabolites exchange at varying oxygen uptake rates in *Sint-GEM*. Exchange flux (in mmol/gDW h) of galactitol (blue solid), ethanol (dotted light blue), and CO_2_ (black dash line) shown on left y-axis, with specific growth rate (solid red line) on the right y-axis, are plotted against oxygen uptake rate (in mmol/gDW h). E) Cell density and extracellular metabolite concentration over time for wild-type *S. intermedia* grown on batch reactors with low aeration levels (2% O_2_). Biomass (OD, represented with diamonds), extracellular concentration of galactose (squares), galactitol (circles) and ethanol (triangles). F) Percentage of O_2_ (aqua) and CO_2_ (purple) measured at the air outlet along the cultivation time in batch reactors with low aeration levels. Vertical grey dotted lines delimit the time span with the highest measured growth and galactose consumption rates.

The observed aerobic galactitol production on galactose, contrasted with its absence in *Sint-GEM* predictions under galactose-limited conditions, suggests that galactitol secretion may function as an overflow metabolism mechanism. This secretion could result from incomplete respiration of incoming galactose. Additionally, regenerated NAD^+^ coupled to galactitol overflow may be consumed by a high glycolytic flux, analogous to NAD^+^ regeneration by ethanol formation observed in *S. cerevisiae* [49]. This hypothesis could be thoroughly explored with the construction of an enzyme-constrained version of *Sint-GEM*, as this has been a useful modelling approach to shed light on overflow metabolism in other organisms across kingdoms of life [50–53].

Additionally, D-tagatose was detected in trace amounts at the end of batch reactor cultivations of wild-type *S. intermedia* using either galactose or lactose-rich cheese whey permeate as carbon sources (Fig. S6). In the galactose case, sorbitol was also detected in trace amounts. These observations support the predictions from random sampling and show that although secretion of these byproducts may be rare, it does happen (Table 1).

#### Oxygen-limiting conditions

Random sampling and FBA predictions of the *Sint-GEM* solution space both showed that secretion of ethanol is coupled to oxygen-limited cellular growth on galactose and lactose for most of the tested simulations (>94% of the cases) (Table 1) (Fig. 6d). Additionally, almost 60% of the simulations using galactose predicted a coupling between ethanol and galactitol secretion under oxygen limitation. This coupling was also observed experimentally in the reactor cultivations using galactose and low oxygen level (Fig. 6e, f). On the other hand, secretion of galactitol by *S. intermedia* in aerobic reactor cultivations (Fig. 6b, e), suggests that its production might not be strictly dictated by oxygen availability.

Production of ethanol under low oxygen conditions is a widely observed phenotype among budding yeasts and has been described as mechanism for NAD^+^ regeneration (bioreaction engineering principles, pp. 26). This trend was predicted by *Sint-GEM* to be even more pronounced for the growth on lactose cases (100% of simulations) (Table 1). A detailed exploration of flux distributions indicated that, in these scenarios, ethanol production serves to balance the NADH generated from the glycolytic oxidation of the glucose moiety in lactose. Parallelly, galactitol secretion balances the additional NADH that is generated from the galactose moiety, mostly catabolized by the Leloir pathway down to glucose-1-phosphate and further integrated into glycolysis (Suppl. Table 6).

#### Nitrogen-limiting conditions

Further exploration of cellular growth by constraining nitrogen availability on *Sint-GEM* indicated that galactitol secretion is favored by nitrogen-limited growth on galactose and even more on lactose (Table 1). We tested if nitrogen levels in the culture media could affect the secreted galactitol by growing the wild-type strain in bioreactors with only 0.001% ammonium sulphate (referred to as low nitrogen) as opposed to the normal concentration in the Verduyn minimal media (0.5%, referred to hereafter as high nitrogen). Strikingly, galactitol yields were around 3-fold higher in the low compared to the high nitrogen condition, although growth was significantly hampered (Fig. S4).

Overall, the results on quantitative fermentation, supported by model predictions, point out that extracellular accumulation of ethanol and galactitol together are more probable under oxygen limitations. However, galactitol secretion alone was observed in aerobic cultivation on galactose pointing towards its role as an overflow metabolite. Furthermore, increase in galactitol secretion was observed under nitrogen-limited conditions. Notably, these results also show that galactitol accumulation is not an exclusive trait of the *galΔ* mutant but also take place in the wild-type strain.

### Deletion mutants and gene expression suggest regulatory crosstalk between Leloir and oxidoreductive pathways

The network topology of *Sint-GEM*, featuring the hypothesized oxidoreductive pathway, prompted the construction of deletion mutants to verify the involvement and function of specific enzymes within this pathway. To this end, we assessed growth on galactitol and sorbitol for wild-type and deletion mutants of the most upregulated genes in the oxidoreductive pathway *(xyl1_2Δ, lad1Δ, lxr4Δ* and *xyl2Δ*) as well as individual Leloir genes (*gal1Δ, gal7Δ* and *gal10Δ*) (Fig. 7). Interestingly, *gal1Δ* and *gal7Δ* deletion mutants in the Leloir pathway and *xyl1_2Δ* in the oxidoreductive pathway displayed severe growth defects on galactitol but not on sorbitol. In contrast, *xyl2Δ* showed an extended lag phase on sorbitol as compared to the wild-type strain, suggesting that the predicted oxidoreductive pathway structure rightfully accounts for the role of Xyl2 as a possible sorbitol dehydrogenase in *S. intermedia*, although further verification is needed. The oxidoreductive pathway deletion mutants *lad1Δ* and *lxr4Δ* did not show any growth defects on either of the carbon sources tested, in fact they seem to grow better than wildtype on galactitol. Results from orthology searches revealed that these genes have no predicted paralogs in *S. intermedia*. Thus, absence of growth-impaired phenotypes may be a consequence of a high degree of substrate promiscuity of oxidoreductive enzymes [38, 54], meaning that other oxidoreductases can compensate for the function of the enzymes encoded by these genes, despite a lack of sequence similarity. Additionally, carbon flow from galactitol may also be routed through alternative pathways that are not represented in the *Sint-GEM.* This phenomenon could be further explored through metabolomic analysis and identification of pathway intermediates in different deletion mutants growing on galactitol.

**Figure 7.**
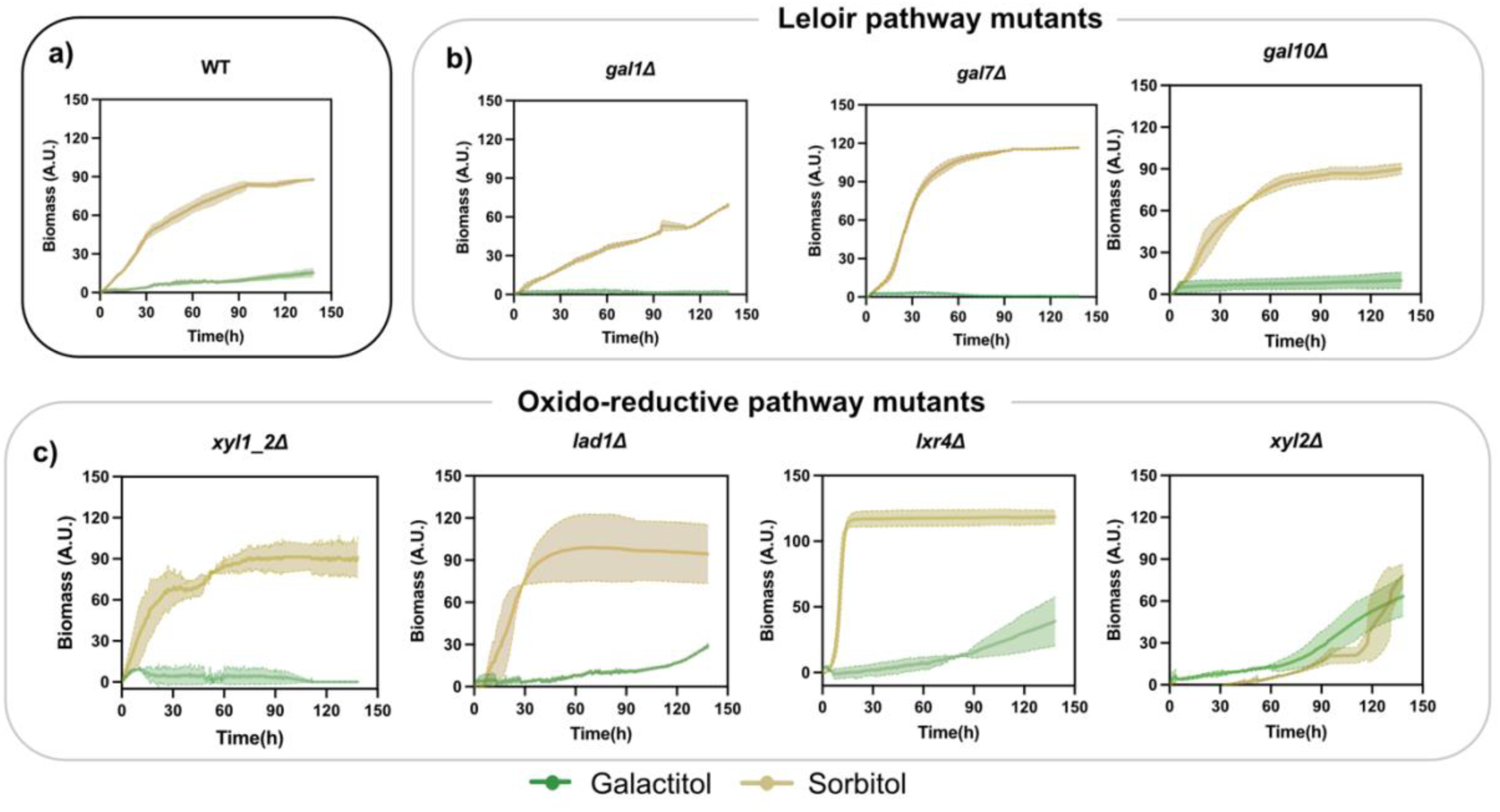
Growth curves for wild-type and deletion mutants. A) Wild-type, b) Leloir pathway mutants and c) hypothesized oxidoreductive pathway mutants in minimal media containing 2% galactitol or 2% sorbitol as carbon source. Graph indicates Time (h) on x-axis plotted against Green Values (A.U) as determined in a 96 well plate in the growth profiler equipment.

From these results, we can conclude that a purely metabolic network perspective where regulatory mechanisms are ignored, cannot fully account for the observed growth phenotypes across most of these deletion mutants. This motivated a correlation analysis on gene expression levels (RNAseq) across multiple carbon sources (glucose, lactose, galactose, cellobiose and xylose). We found that the expression of *XYL1_2* was tightly correlated to expression patterns of all genes encoding for the Leloir pathway, as well as the lactose transporter *LAC4* and *LAC9* genes (p-value ≤0.01). This result that can be attributed to *XYL1_2* being identified as part of the *GALLAC* cluster [18] (Fig. S5a-b). Moreover, a significant positive correlation was computed between *LAD1*, *XYL1_2*, and *XYL1_3* as part of the oxidoreductive pathway, and between *LAD1* and all genes in the Leloir pathway. *LXR4* showed significant correlation to *XYL1_2, LAD1, GAL1_2* and *LAC9*, while *XYL2*, encoding for the final step in the oxidoreductive pathway, was found to be significantly correlated only with *XYL1_2* expression. These results suggest a complex network of co-expression and/or co-regulation of genes within both the Leloir and oxidoreductive pathways, as well as crosstalk among genes of both pathways. The observed gene expression correlation, along with the unexpected growth defect on galactitol of the *gal1Δ*, *gal7Δ*, and *xyl1_2Δ* mutants, suggests possible regulatory effects on downstream genes in the oxidoreductive pathway.

### *Sungouiella intermedia* has cell factory potential for production of galactitol and tagatose

Finally, we set out to combine *Sint-GEM* modelling and experimental analysis to optimize the bioprocess design for the *S. intermedia galΔ* strain as cell factory on lactose-rich feedstocks. We first conducted flux sampling to identify the conditions under which this mutant strain secretes galactitol. Simulations on lactose indicate that ethanol and galactitol are the predominant secreted byproducts, accounting for 39.4% and 30.9% of the simulations, respectively. Notably, they are often predicted to be co-secreted, occurring in 26.3% of cases, particularly under oxygen-limited conditions. On the other side, random sampling of the *galΔ* model under carbon-limited conditions showed very few cases of ethanol secretion (0.14%). For experimental verification, we grew the *galΔ* strain in minimal medium with lactose in both aerobic and low-oxygen (2%) conditions in batch reactors. Extracellular galactitol was detected in both conditions (4.8 and 1.2 g/L, respectively) and was accompanied by secreted ethanol (1.1 g/L) in the low-oxygen case (Fig. S7). These results provide important guidance for choosing an oxygen-rich bioprocess design when applying *S. intermedia galΔ* as a production host for conversion of lactose-rich whey into bioproducts.

We also assessed the bioproduction process design, with the goal of determining and improving the titer, rate and yield of galactitol production. Having observed that the *galΔ* strain exhibits growth after an extended lag phase (∼120 hours) in minimal media with 2% lactose after being precultured in glucose-containing media and recognizing that galactitol is a growth-coupled metabolite, we hypothesized that preculturing in lactose would enhance productivity. Indeed, preculturing the *galΔ* strain in medium containing lactose instead of glucose reduced the lag phase to 48 h, thus improving the overall productivity in shake flasks from 0.015 g galactitol/L h^-1^ to 0.042 g galactitol/L h^-1^ and reaching a titer of 7.2 g/L and a yield of 0.45 g galactitol produced per g lactose consumed (85% of the stoichiometric theoretical maximum) (Fig. 8a). Following the same culturing sequence, we assessed whether the final titer of galactitol could be improved by growing the strain in minimal media containing 5% lactose (instead of the original 2%) and using a 1L bioreactor to ensure fully aerobic conditions as suggested by *Sint-GEM* simulations and experimental analyses. Indeed, we observed a noteworthy increase in titer and rate of 21 g/L and 0.11 g/Lh^-1^, respectively (Fig. 8b), while the final yield of 0.42 g of galactitol produced per g lactose consumed was not improved compared to the previous experimental setup. No extracellular ethanol could be detected, which was in accordance with model predictions.

**Figure 8:**
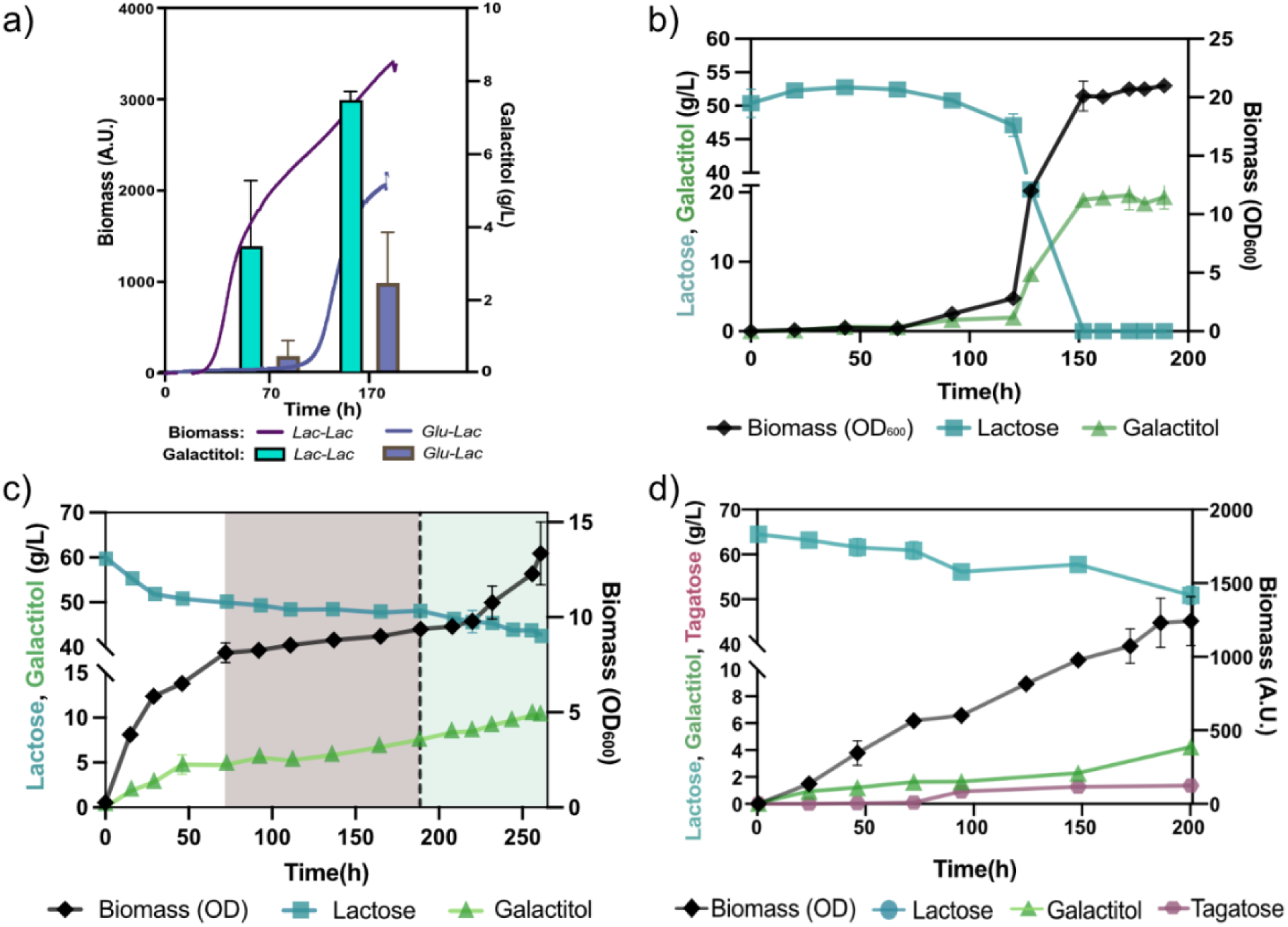
Production of galactitol and tagatose using *S. intermedia* as a cell factory. A) Galactitol titers in g/L plotted over time (h) as bar plot for two culturing schemes, Glu-Lac (green): The strain is grown on solid media containing glucose as carbon source, precultured into liquid media containing glucose and final culture is in minimal media containing lactose and Lac-Lac (purple): The strain is grown on solid media containing lactose as c-source, precultured into liquid media containing glucose as c-source and final culture is in minimal media containing lactose. Error bars represent standard deviations for two biological replicates. B) Galactitol production in 1L batch reactor containing minimal media with 5% lactose. Biomass in cell dry weight (empty circle) is plotted on the right y-axis and lactose (filled circle), glucose (empty square), galactose (empty triangle) and galactitol (empty reverse triangle) is plotted on left y-axis, against time (h) on x-axis. Error bars represent standard deviation for two biological replicates. C) Galactitol production in 1L bioreactor containing CWP with starting lactose concentration of 60 g/L. Dotted line depicts the addition of ammonium sulphate to the culture. Shaded regions have been compared for change in galactitol yield, growth rate and lactose uptake rates to determine the effect of ammonium sulphate addition. Biomass (OD) is shown in black and plotted on right y-axis, lactose (blue) and galactitol (green) plotted on left y-axis, against time(h) on x-axis. Error bars represent standard deviation for two biological replicates. D) Tagatose production by *galΔade1::GDH* strain in shake flasks containing CWP with a starting lactose concentration of 60g/L. Biomass (OD) is shown in red and plotted on right y-axis, lactose (green) and galactitol (light-green) plotted on left y-axis, against time(h) on x-axis. Error bars represent standard deviation for two biological replicates.

To assess the applicability of the *S. intermedia galΔ* strain for conversion of a real industrial side-stream, we cultivated the strain in cheese whey permeate (CWP) containing 6% lactose in fully aerated 1L bioreactors. In contrast to the relatively high titers observed in cultures in minimal media with 5% lactose, on CWP, the strain only attained 5g/L galactitol as the highest titer after 150h (Fig. 8d). However, galactitol yield remained high (0.54 g galactitol produced per g lactose consumed) during the whole duration of fermentation, which could be attributed to low nitrogen availability in CWP [55], in accordance with the results from our previous cultivations in low nitrogen. To confirm if nitrogen availability influences the galactitol yield under these conditions, we decided to add ammonium sulphate to a final concentration of 1g/L, and compared fermentation metrics before and after the addition of ammonium sulphate (gray and green shaded area in Figure 8c, respectively). After addition, we observed an immediate increase in growth rate (0.01 v 0.07 h^-1^) coupled with faster lactose uptake rate (0.01 vs 0.07 g/L h^-1^) and higher galactitol production rate (0.024 vs 0.035 g/L h^-1^). The final titer increased to 10.2 g/L, while the yield of galactitol production remained unchanged. This may be attributed to the relatively low final concentration of ammonium sulphate in the CWP as compared to that in minimal media (0.01 vs 0.05%). Additionally, the duration of the cultivation after addition of ammonium sulphate may not have been sufficient to observe a pronounced difference in titer.

Next, we aimed to engineer *S. intermedia* to produce the natural sweetener tagatose, via galactitol. The tagatose producing strain was constructed using CRISPR-Cas9 mediated genome integration of the galactitol-2-dehydrogenase-encoding gene *RlGDH* from *Rhizobium legumenosarum*, as previously demonstrated by Jin and colleagues [21]. The gene was integrated in the *ADE1* locus of the *S. intermedia galΔ strain,* creating the strain *galΔ, ade1::GDH*. Upon cultivation in CWP, the strain converted parts of the produced galactitol into tagatose (Fig. 8d), resulting in extracellular accumulation of 4.1 g/L galactitol and 1.3 g/L tagatose. The relatively low tagatose titer indicates that expression of galactitol-2-dehydrogenase by *RlGDH* gene needs further optimization to achieve higher conversion to tagatose. Overall, our results demonstrate the applicability of *S. intermedia* strains as a potential cell factory for production of added-value products using CWP as feedstock.

## Conclusions

In this work, we have demonstrated the existence of an oxidoreductive pathway that operates in parallel with the Leloir pathway for galactose catabolism in the yeast *S. intermedia.* Furthermore, we have leveraged the insights gained throughout this study to showcase the potential applications of this yeast as a cell factory.

Here, a draft genome-scale metabolic model was curated using experimental data from bioreactors, including exchange fluxes and biomass composition. This model was further enhanced by incorporating orthology searches across filamentous fungi and RNA-seq data from growth on various carbon sources, resulting in the high-quality metabolic network model *Sint-GEM* that is well-suited for the simulation and quantitative exploration of the metabolic landscape in *S. intermedia*. Analysis of the predicted flux distributions indicated that the galactose oxidoreductive pathway is a crucial component of *S. intermedia*’s lactose and galactose metabolism, allowing it to adapt to various nutrient limitations (carbon, oxygen, and/or nitrogen). This pathway may also serve as a mechanism for managing excess carbon, potentially functioning as part of an overflow metabolism strategy.

Moreover, simulations predicted that versatile use of this pathway can lead to secretion of diverse byproducts which, at least in the case of galactitol, can be assimilated back into the cell as a carbon source under situations of nutrient scarcity. As galactitol is a product known to be toxic to other yeasts [56], this predicted and observed phenotype might provide *S. intermedia* cells with an adaptive advantage over other competing species in lactose/galactose rich niches.

Ultimately, the insights derived from the knowledge-matching process of *Sint-GEM* curation, simulation, and experimental analysis were utilized to demonstrate *S. intermedia*’s potential as a cell factory for lactose-rich industrial byproducts. This involved optimizing process conditions to enhance galactitol productivity and introducing a heterologous gene to boost the production of its derivative, tagatose.

## Materials and Methods

### *Sint-GEM* structure

*Sint-GEM* constitutes a catalogue of all enzyme-encoding genes, reactions and metabolites predicted to be part of *S. intermedia* metabolism. Metabolic reactions are represented by a stoichiometric matrix (***S***), which represents reactions in its columns and metabolites in its rows. Coefficients in this matrix represent the stoichiometric coefficient of a metabolite in each reaction it participates. As convention, negative coefficients are used for reaction substrates, while positive values represent reaction products. A two reactions example would take the following form:

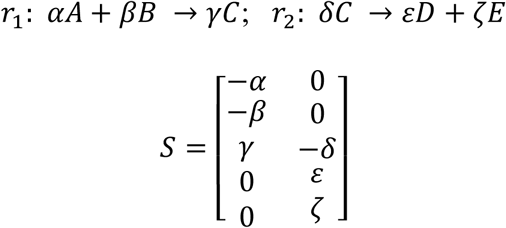

Similarly, relations between reactions and genes are stored in a reaction-gene matrix (***RG***), in which rows represent reactions in the model and columns the genes. If a gene product (enzyme) catalyzes a given reaction, then the corresponding coefficient in the ***RG*** matrix is assigned a value of 1. Rows with multiple non-zero elements indicate reactions with isoenzymes (i.e., different enzymes with equivalent catalytic activity) or reactions catalyzed by enzyme complexes, columns with multiple non-zero coefficients indicate promiscuous enzymes (i. e., enzymes with catalytic activity for multiple substrates and reactions). To differentiate between enzyme complexes or reactions with isoenzymes, an additional field relating reactions to genes is added to the model, *grRules*, indicating the gene(s) encoding for all enzymes related to each reaction. Rules of the form *g*_*i*_ *or g*_*j*_ indicate reactions with isoenzymes. Rules of the form (*g*_*i*_ *and g*_*j*_) indicate reactions catalyzed by enzyme complexes, in this case composed by subunits *i* and *j*.

### Standardization of *Sint-GEM*

A draft model reconstruction, based on yeast8 consensus metabolic model for *S. cerevisiae* and orthology searches across 332 budding yeasts and 11 filamentous fungi annotated genomes from a previous study was downloaded from: https://github.com/SysBioChalmers/Yeast-Species-GEMs/blob/main/Reconstruction_script/ModelFiles/xml/Candida_intermedia.xml.

Missing steps in the galactose oxidoreductive pathway were inferred from orthology searches between *S. intermedia* genome and those of *Aspergillus niger, Aspergillus nidulans* and *Trichoderma reesei* using OrthoFinder v.2.5.4. Missing transport steps for lactose and galactitol were added based on observed phenotypes.

Biomass composition was modified in the model in accordance with total protein and total lipids measurements from *S. intermedia* cultures. Amino acid composition for the global protein synthesis pseudoreaction in the model was modified according to amino acid ratios in the protein sequence of *S. intermedia* CBS 141442 strain. Growth-associated and non-growth associated maintenance ATP expenditure, as well as the P/O ratio in the electron transport chain were fitted using RMSE minimization in prediction of experimental values for glucose and oxygen uptakes, and carbon dioxide production from glucose-limited chemostats at 0.05 and 0.1 h^-1^ dilution rate.

Curation, parameter fitting, and model standardization were tracked using the git version control system and adhering to standard-GEM guidelines^14^, thus facilitating reproducibility, traceability and further development of the resulting GEM^15^. All model files and curation scripts are publicly available at: https://github.com/SysBioChalmers/cint-GEM.

### FBA simulations for optimal growth

Cellular growth simulations under a given nutrient limitation were performed using the flux balance analysis method, which assumes steady state for the intracellular concentrations of metabolites, and phenotype prediction by optimization of a cellular objective, thus yielding the following linear programing problem:

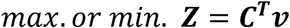

Subject to

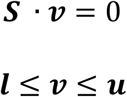

Where *Z* is the optimal value of the objective function; *C*^*T*^ is a row vector of coefficients of the objective function for each element in *v*; *v* is the vector of intracellular reaction fluxes, in units of mmol/gDw h; *S* is the stoichiometric matrix described above; *l* and *u* are vectors of lower and upper bounds, respectively, for the reaction fluxes in *v*. Cellular growth was simulated maximizing the biomass production pseudoreaction in *Sint-GEM* (r_4041), setting an arbitrarily low value of 1000 mmol/gDw h as lower bound for the exchange reactions of components in minimal essential media, and a decreasing value from 0 to −1 mmol/gDw h as lower bounds for the galactose/lactose and oxygen exchange reactions, depending on the simulated limiting condition. All exchange reactions were constrained by an upper bound of 1000 mmol/gDw h to allow secretion of metabolites. The RAVEN toolbox v2.8.7was used for all FBA simulations, with Gurobi v.10 as numerical solver and MATLAB R2022b.

### Random sampling of *Sint-GEM* solution space

A series of 10,000 FBA simulations with randomly generated coefficient vector (*C*^*T*^) for the objective function for carbon-, oxygen- and nitrogen-limited conditions under lactose and galactose as sole carbon sources. For each case an initial growth maximization with a unit carbon source uptake rate as a constraint was simulated, as described above, to obtain *v*^∗^_growth_ which is then used to set a lower bound in the growth pseudoreaction as 0.5 ∗ *v*^∗^*_grouth_* (suboptimal growth rate). A lower bound of −1 mmol/gDw h is set as lower bound for the exchange reactions of galactose, lactose exchange in the carbon limited cases, oxygen exchange for oxygen-limited condition, and ammonium for the nitrogen-limited cases. When carbon is not the limiting element, then the lower bound for the galactose or lactose exchange reaction was set to −1000 mmol/gDw h. Minimal media components uptake and exchange metabolites secretion was allowed as described above. All random sampling simulations were run with the *randomSampling.m* function in the RAVEN toolbox.

### Analysis of metabolite turnover distributions

All flux distributions simulated by random sampling were converted to metabolite turnover distributions, in which for each metabolite turnover (τ_*i*_)

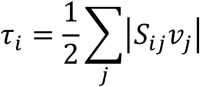

This quantity represents the total rate of transformation of metabolite *i* across all reactions in which it partakes (as substrate or product). For each simulated limiting condition, a matrix containing each of the computed metabolite turnover distributions for the respective 10,000 simulations as columns. Dimensionality reduction by principal component analysis (PCA) was performed on such matrix for each of the simulated conditions using the *prcomp* function in R v4.4.1. The top metabolites contributing to the mapping of samples (metabolite turnover distributions of all simulations) onto the first two principal components were obtained by identifying all metabolites with a weight coefficient on the same order of magnitude as the maximum coefficient for each PC (the minimum for the negative directions in PCs).

### Gene expression correlation analysis

For all genes included in *Sint-GEM,* normalized RNA expression values across conditions (glucose, lactose, galactose, cellobiose and xylose as sole carbon sources) were obtained from the RNAseq dataset. Pearson correlation coefficient and t-student statistical test was computed for all possible pairs of gene expression vectors using the *corrcoef* function in MATLAB R2022B.

### Determination of biomass composition

To determine biomass composition, *S. intermedia* was cultivated in a carbon-limited chemostat. An overnight culture was prepared in minimal Verduyn media [57] containing 2% glucose or lactose. It was then inoculated into a controlled stirred 1L bioreactor vessels (DASGIP, Eppendorf, Hamburg, Germany) containing 500 mL minimal Verduyn media with 2% glucose or lactose, respectively. Reactor conditions were maintained as: Temp = 30 °C; pH = 5.5 (maintained with 2M Potassium Hydroxide); Aeration = 1 Vessel Volume per Minute; stirring = 300 rpm. The cultures were allowed to grow until stationary phase before turning on dilution of 0.1 h^-1^ and 0.05 h^-1^. After the cultures had reached steady state (as determined by CO_2_out and cell dry weight measured at an interval of at least 12 h), cells were harvested for measurement cell dry weight, total proteins and lipids as outlined below.

The original Yeast 8 model’s biomass reaction was used as the starting point for the biomass reaction in draft *Sint-GEM*. Experimentally determined specifications from *S. cerevisiae* already present in *Sint-GEM* were taken; specifically amino acid, inorganic compounds (i.e., phosphate, sulfate, and metal ions), and cell wall compositions, as corresponding data for *S. intermedia* CBS 141442 were not available. Additional data adopted from the yeast 8 model were Genome GC content, relative abundance of acyl groups and free fatty acids in lipids, lipid subspecies composition (e.g., phosphatidylinositol, phosphatidylcholine, phosphatidylethanolamine, and phosphatidylserine composition). We used total protein and lipid composition, generated experimentally, to fine-tune the metabolite coefficients in the biomass reaction in *Sint-GEM*. Metabolite coefficients associated with growth-associated ATP maintenance were updated in *Sint-GEM* using experimentally determined rates from bioreactor cultures (Suppl. Table 5).

#### Total proteins

Protein was measured using the Lowry method. Briefly, 20 mL culture was pelleted by centrifugation at 10,000 x g for 5 min, washed and resuspended in 10 ml. An aliquot of 2 ml from this is mixed with 1 ml of 3M NaOH. Cell suspension was incubated at 100 °C for exactly 10 min and then immediately cooled on ice. An aliquot of 0.45 ml of the sample was taken in a clean tube, to which 0.45 ml 1M NaOH and 0.3 ml of Copper Sulphate (2.5% w/v) was added. The solution was mixed well and incubated at room temperature for 5 min, following which, the mix was centrifuged at 10,000 x g for 5 mins. Absorbance was measured of the clear supernatant at 510 nm. For calibration curve, Bovine Serum Albumin (Thermo), at a starting concentration of 5mg/ml was used.

#### Total lipids

Dry biomass was weighed in glass tubes (100 mg) and resuspended in Chlorophorm Methanol (2:1) solution and heated to 80 °C. Tubes were flushed with nitrogen and dried overnight to measure the total lipid in each sample. Finally, tubes were weighed to determine the total lipid content per g of biomass.

### RNAseq and data analysis

Transcriptomics data from RNA sequencing was adopted from previous work [15]. In summary, *S. intermedia* CBS 141442 was grown in controlled stirred 1-L bioreactor vessels (DASGIP, Eppendorf, Hamburg, Germany) containing 500 mL minimal Verduyn media with 2% Glucose, Galactose or Lactose. Reactor conditions were maintained as: Temp = 30 °C; pH = 5.5 (maintained with 2 M Potassium Hydroxide); Aeration = 1 Vessel Volume per Minute; stirring = 300 rpm. Cells were harvested (10 mL) when the dissolved oxygen of the culture was 35 to 40% (v/v) and subsequently snap-frozen and stored in −80 °C for extraction using TRIzol-chloroform method. RNA samples were analyzed in a TapeStation (Agilent, Santa Clara, CA, USA) and sequencing was performed using HiSeq 2500 system (Illumina Inc.—San Diego, CA, USA) as described previously [15].

Gene counts were normalized with weighted trimmed mean of M-values using the calcNormFactor function from the package edgeR [58]. Limma package was used to transform and make data suitable for linear modelling [59]. The estimated p-values were corrected for multiple testing with the Benjamini-Hochberg procedure, and genes were considered significant if the adjusted p-values were lower than 0.01. The raw counts were filtered such that genes with counts per million > 3.84 in at least 12% (5/43) of the samples were retained. The function ‘varianceStabilizingTransformation()’ from R package ‘DESeq2’69 was used to convert raw counts to variance-stabilized-counts (VST). Expression data for *S. intermedia* on galactose and lactose was normalized using glucose as control condition. The RNA seq datasets are available in the European Nucleotide Archive (ENA) with the accession number E-MTAB-6670 [18].

### Molecular techniques and amplification of plasmid

Golden Gate cloning Gate cloning [60] was used to construct the galactitol-2-dehydrogenase integration fragment consisting of the *SiPET9* promoter driving the *Rl*GDH gene with *CiACT1* terminator, in the BsmBI cloning site of the plasmid *pGGAselect* ordered from Addgene (https://www.addgene.org/195714/). PCR amplification of the integration fragment from the plasmid was performed using primers with 50bp overhangs homology to up/down-stream of the *SiADE1* gene. Integration fragment and the CRISPR-Cas9 plasmid (constructed for gene editing in *S. intermedia*) containing the guide RNA targeting the *SiADE1* gene were used to construct the *galΔade1*::*GDH* strain. Plasmid and integration fragment were transformed and selected based on resistance conferred to nourseothricin (25 µg/ml) using a previously described protocol [17].

### *E. coli* transformation

For amplification of plasmids, *E. coli* DH5α cells were transformed using heat-shock method, followed by selection of ampicillin (100 µg/ml). Transformants screened by colony PCR were subsequently grown on LB medium (1 % tryptone, 1 % NaCl and 0.5 % yeast extract) containing chloramphenicol (25 μg/mL) or ampicillin (100 μg/mL) for plasmid selection.

### Yeast transformation

Targeted gene deletions *galΔ*, *lad1Δ*, *lxr4Δ*, *xyl2Δ* was performed using the split-marker technique and transformation through electroporation, as described in detail in our previous work [17]. After transformation, cells were plated on YPD agar containing 200 μg/ml nourseothricin to select for integration and expression of the *Ca*NAT1 selection marker.

### Screening for transformants

Nourseothricin resistant transformants were screened for the targeted deletion or insertion using colony PCR. Single colonies were picked and transferred to 200 μl PCR tubes containing 20 μl milliQ using sterile toothpicks. The PCR tubes were placed in a thermocycler and heated to 90°C for 10 mins followed by subsequent cooling to 12 °C. For each colony picked, 2 μl of the heated colony material was used as template for PCR using PHIRE II polymerase. Screening for transformants using PCR was performed by employing primer pairs with one primer binding to the genome outside the homology flank of the integration fragment and one on the marker gene. A third primer binding to the target gene was used added to the mix, to identify transformants with incorrect integration.

### Growth Characterization

#### Growth profiler

Wild-type and mutant strains of *S. intermedia* were cultivated in 96-well plate format using the Growth Profiler 960 (Enzyscreen, The Netherlands) to follow growth over time. Strains were precultured at 30 °C, 180 rpm overnight in minimal Verduyn media with different carbon sources. Precultured cells were then inoculated in 200 µL minimal media supplemented with 2% carbon source (1% in case of galactitol) to a starting OD600 = 0.1. ‘Green Values’ (GV) measured by the Growth Profiler 960 (EnzyScreen, The Netherlands) correspond to growth based on pixel counts, and GV changes were recorded every 30 min for 72 h at 30 °C and 250 rpm.

#### Cell growth quantifier (CGQ)

Growth characterization was also performed in 100 ml shake flasks containing 25ml cultures using the CGQ system (aquila biolabs GmbH, Germany) [61]. Wild-type and mutant strains were precultured at 30 °C, 200 rpm overnight in Verduyn minimal medium containing 2% glucose (w/v), and thereafter inoculated in 25ml of Verduyn minimal medium supplemented with 2% carbon source (galactose or lactose) in 100ml shake flasks to a starting OD_600_ of 0.1. Growth was quantified as “Scatter values” by the aquila system-based on the scatter light detected by the sensors.

#### Growth in bioreactors

In addition to characterization of the wild-type strain in carbon limited chemostats, *S. intermedia* strains (wild-type (*S. intermedia* CBS141442), *galΔ* and *galΔ::RlGDH*) were cultured in stirred 1L bioreactors (DASGIP, Eppendorf, Germany) containing 500 mL of either minimal Verduyn media containing 5% lactose or industrial cheese whey permeate (WheyCo Inc.). Batch reactor runs were performed at 30 °C, pH = 5.5 (maintained with 2M Potassium Hydroxide) and aeration coupled with stirring was set to cascade mode to maintain dissolved oxygen at 30% as lowest value. Cells were harvested for biomass measurement and metabolite analysis using HPLC at regular intervals. Cultivation of *galΔ::RlGDH* strain was supplemented with adenine (40 mg/L).

### Metabolite concentrations using HPLC

Changes in metabolites such as lactose, galactose, glucose and galactitol were quantified using Jasco high-performance liquid chromatography (HPLC) system (Jasco Corporation, Japan) equipped with an RID-10A refractive index detector, a Rezex RCM Monosaccharide Ca(^+2^) (8%) column (Phenomenex Inc., USA) with MilliQ Water (Merck MilliPore, USA) as the mobile phase and eluted at a flow rate of 0.6 mL/min at 80 °C. Additionally, detection and quantification of organic acids and ethanol was performed using Rezex-Organic Acid column (Phenomenex Inc., USA) with H_2_SO_4_ as mobile phase and eluted at a flow rate of 0.8 mL/min at 50 °C. For the detection of Tagatose and Sorbose, Dionex ICS-5000+ (ThermoFisher, USA) was applied using the monosaccharide separation column, Dionex™ CarboPac™ PA1 (ThermoFisher, USA). Separation was performed using MilliQ Water as the mobile phase and eluted at a flow rate of 1 mL/min at 70 °C. Prior to analysis, culture samples were pelleted, and the supernatant was passed through 0.22 μm polyethersulfone syringe filter. Chromatogram peaks were treated and integrated using the software ChromNav v2.0 (Jasco HPLC) or Chromeleon v6.8 (Dionex IC).

## Acknowledgements

The authors would like to thank the following members of the Division of Industrial Biotechnology at: Fabio Caputo and Yi-Hsuan Li for assistance with analytics, Vijay Raghavendran, PhD., Marta Mohedano, PhD., and Bruna Ferreira, for their guidance and help with bioreactor cultivations, and Marta Parmigiani for help with molecular work.

## Author contributions

K.V.R.P., I.D. and C.G conceptualized the project; K.V.R.P., I.D., and C.G. designed the methodology; K.V.R.P., I.D., H.A., A.V.R. and C.G. produced and analyzed investigation results; K.V.R.P., I.D., and C.G. wrote original manuscript draft; all authors participated in manuscript review and editing; funding acquisition by C. G. and J. N.

## Supplementary Material

Supplementary Tables 1-6 can be found here: https://github.com/SysBioChalmers/sint-GEM

**Figure S1:**
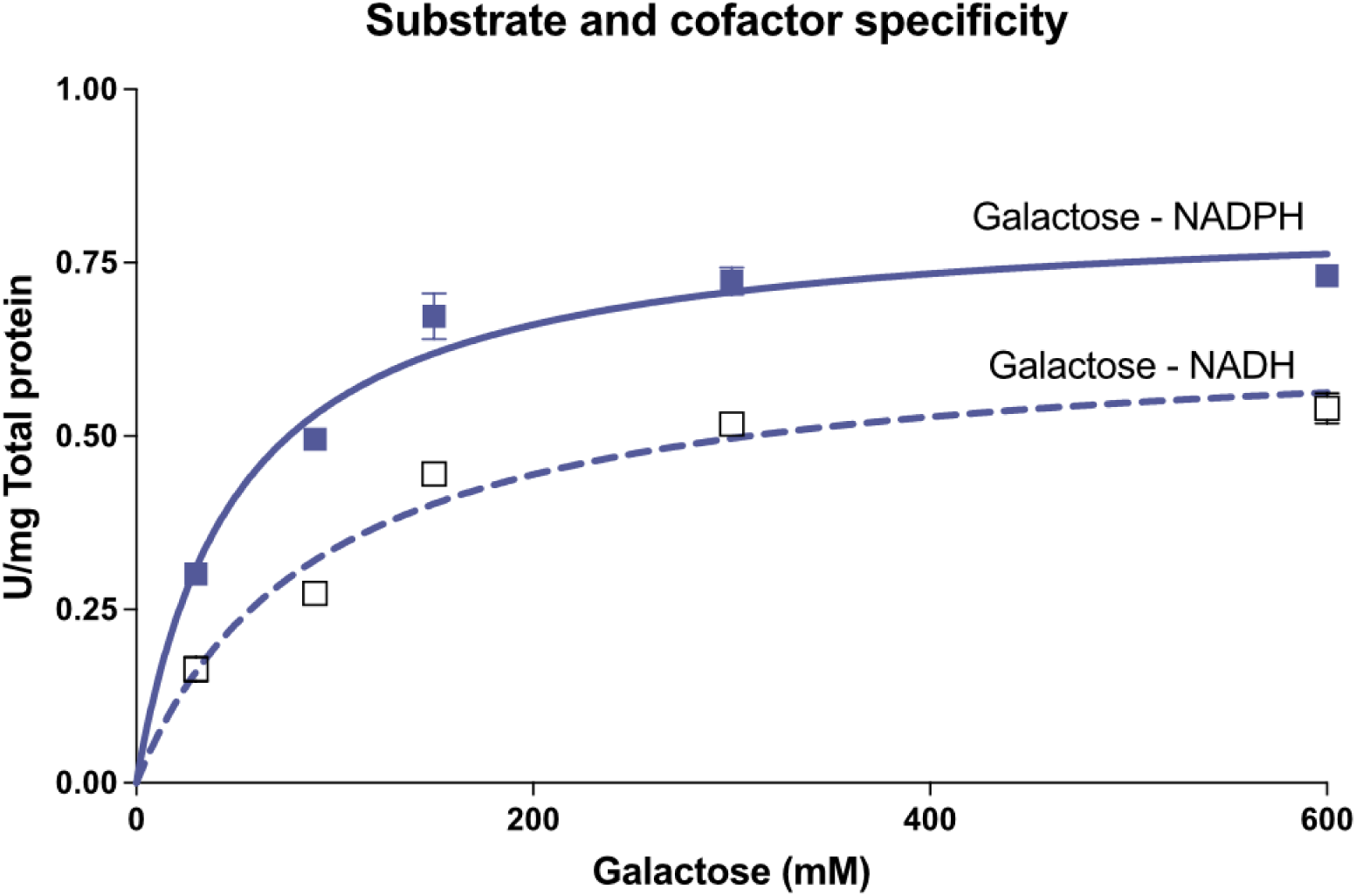
Enzymatic activity for the determination of co-factor specificity of aldose reductase in S. intermedia.

**Figure S2.**
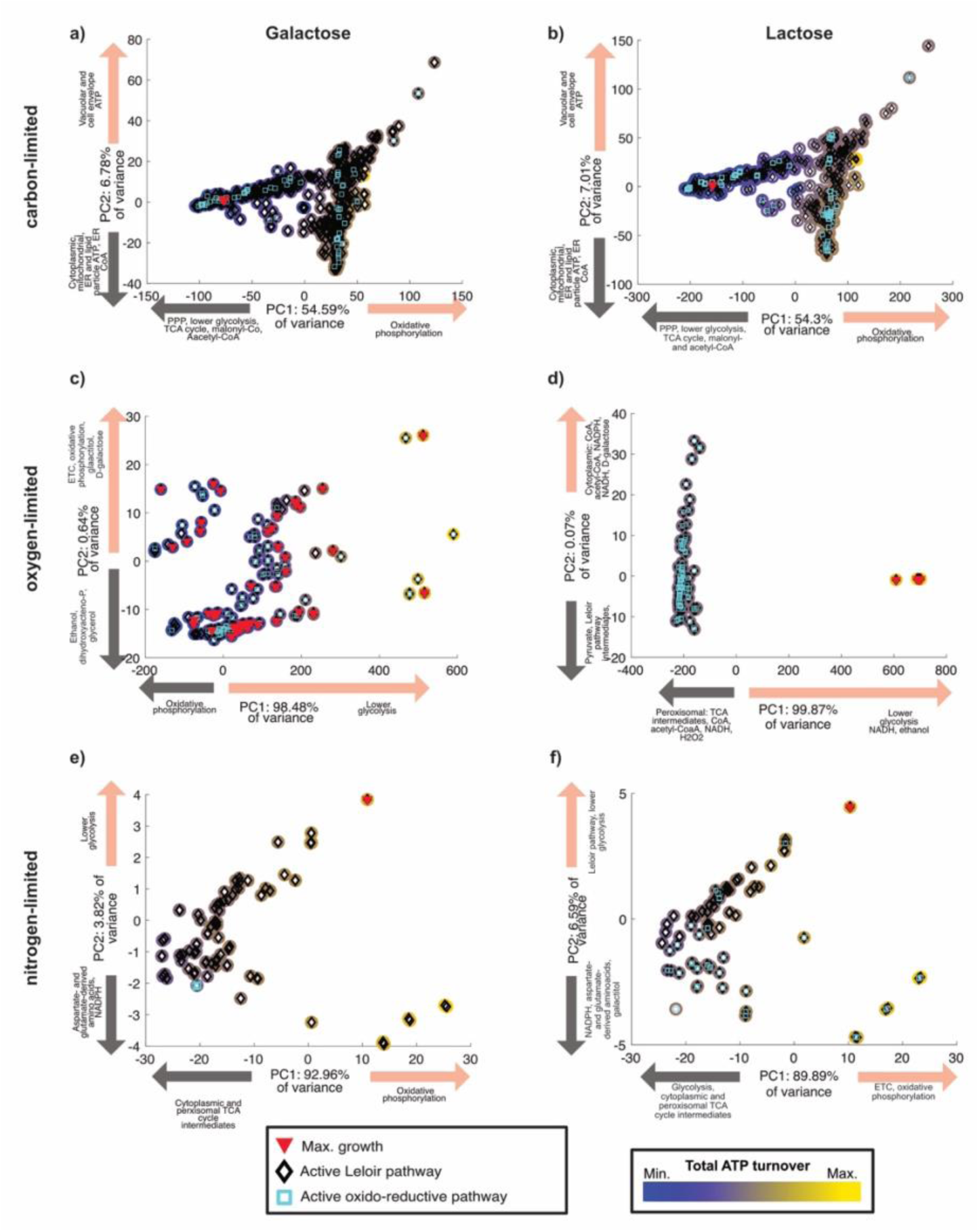
PCA of galactose pathway utilization. 10,000 randomly sampled flux distributions simulations are converted into metabolite turnover distributions for each case. Simulated conditions are growth on galactose under carbon-limitation (a), oxygen-limitation (c), nitrogen-limitation (e); and growth on lactose under carbon-limitation (b), oxygen-limitation (d), nitrogen-limitation (f). Global ATP turnover was mapped onto the color scale for each simulation point (dark blue for minimum ATP turnover – bright yellow for maximum values). Simulations displaying maximum growth rate under the specified constraints are marked with red filled triangles, those in which the Leloir pathway is carrying flux are shown with black non-filled diamonds, and those in which the galactose oxidoreductive pathway carries flux are highlighted with non-filled aqua squares. Metabolites and pathways with the highest contribution to positive and negative values of PC1 and PC2 are displayed along the X and Y axis, respectively.

**Figure S3.**
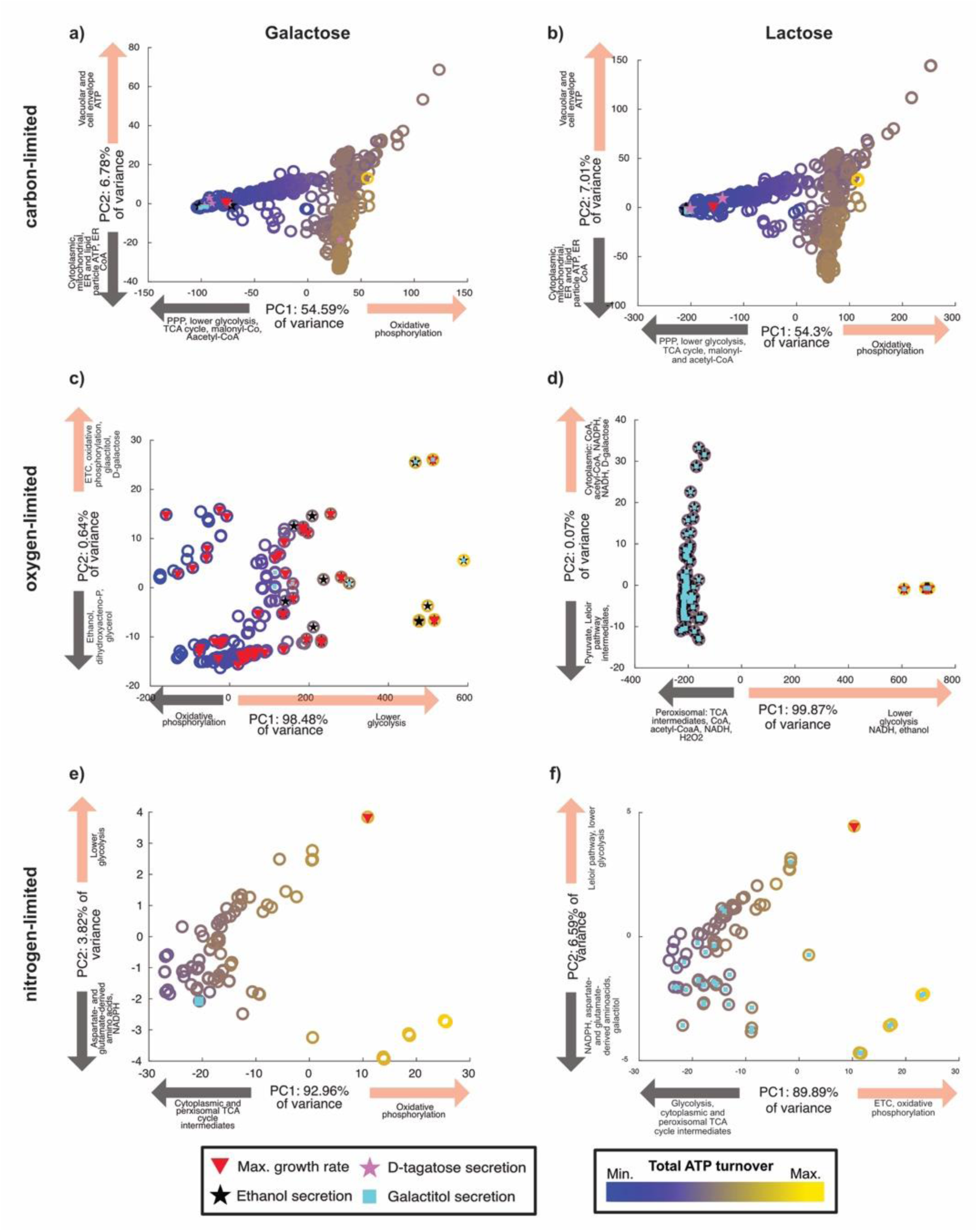
PCA of byproduct secretion. 10,000 randomly sampled flux distributions simulations are converted into metabolite turnover distributions for each case. Simulated conditions are growth on galactose under carbon-limitation (a), oxygen-limitation (c), nitrogen-limitation (e); and growth on lactose under carbon-limitation (b), oxygen-limitation (d), nitrogen-limitation (f). Global ATP turnover was mapped onto the color scale for each simulation point (dark blue for minimum ATP turnover – bright yellow for maximum values). Simulations displaying maximum growth rate under the specified constraints are marked with red triangles, those predicting ethanol secretion are mapped with black pentagrams, D-tagatose secreting simulations with pink pentagrams, and those galactitol secreting simulations with aqua squares. Metabolites and pathways with the highest contribution to positive and negative values of PC1 and PC2 are displayed along the X and Y axis, respectively.

**Fig. S4.**
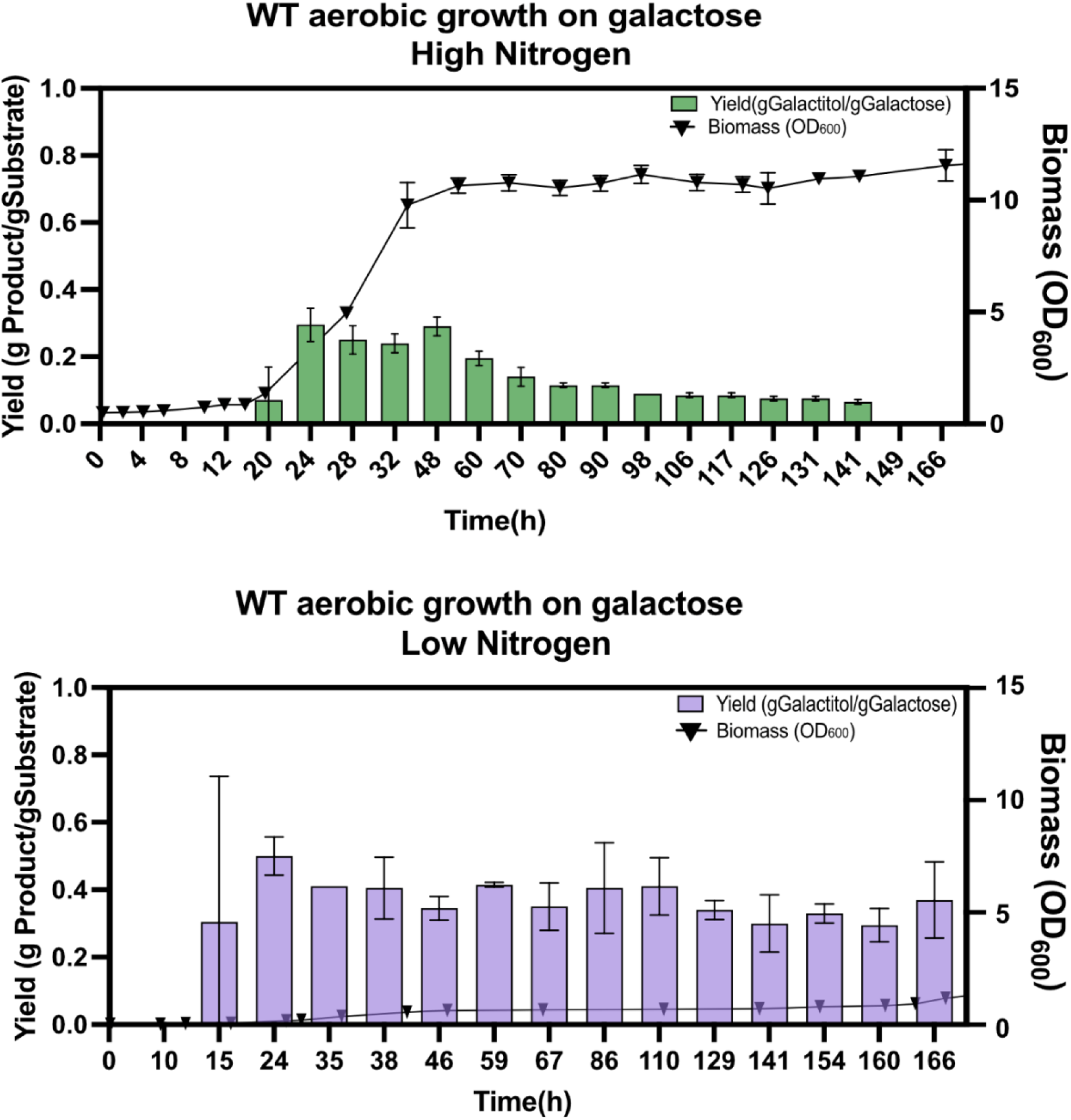
Experimental validation of Sint-GEM results for flux sampling in nitrogen limited condition. Graph depict the yields of galactitol and the biomass of the wild-type strain in minimal media containing two different concentrations of ammonium sulphate (0.5% = high and 0.001% = low).The graphs represent Yield in gGalactitol/gConsumedGalactose on the left y-axis and the biomass (OD600) formation on the right y-axis, plotted against Time(h) on the x-axis.

**Figure S5.**
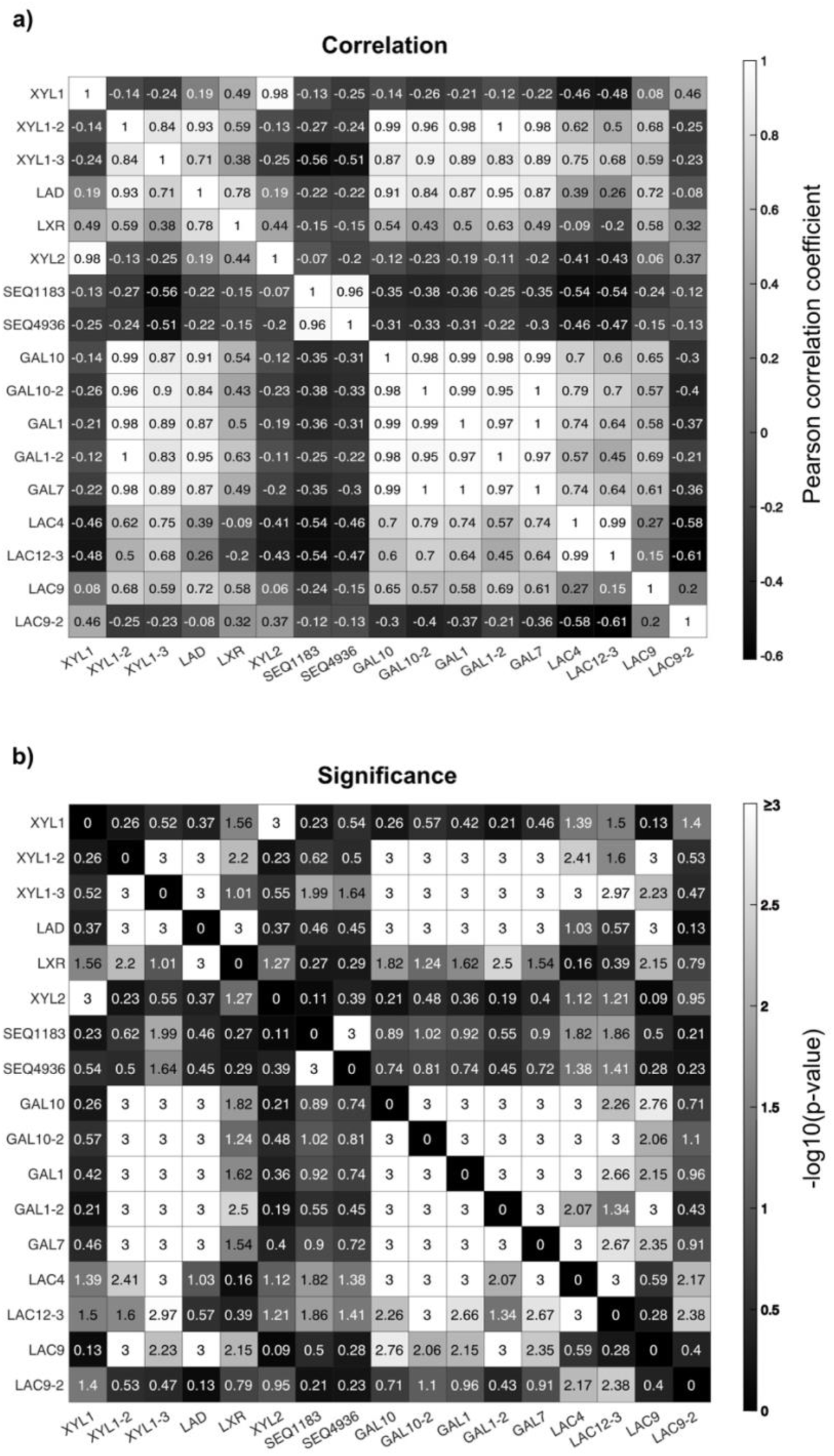
Gene expression correlation analysis. mRNA Expression levels across growth on 5 different carbon sources (glucose, xylose, cellobiose, galactose and lactose) for genes in the oxidoreductive and Leloir galactose metabolizing pathways were compared against each other. a) Pearson correlation coefficients for each pairwise comparison of gene expression vectors are shown with numbers and a grey color scale. b) Significance of linear correlation, expressed as -log10(p-value) is shown in numbers and as a grey color scale. Hypothesis testing was performed using a two-tailed t-test statistic.

**Figure S6:**
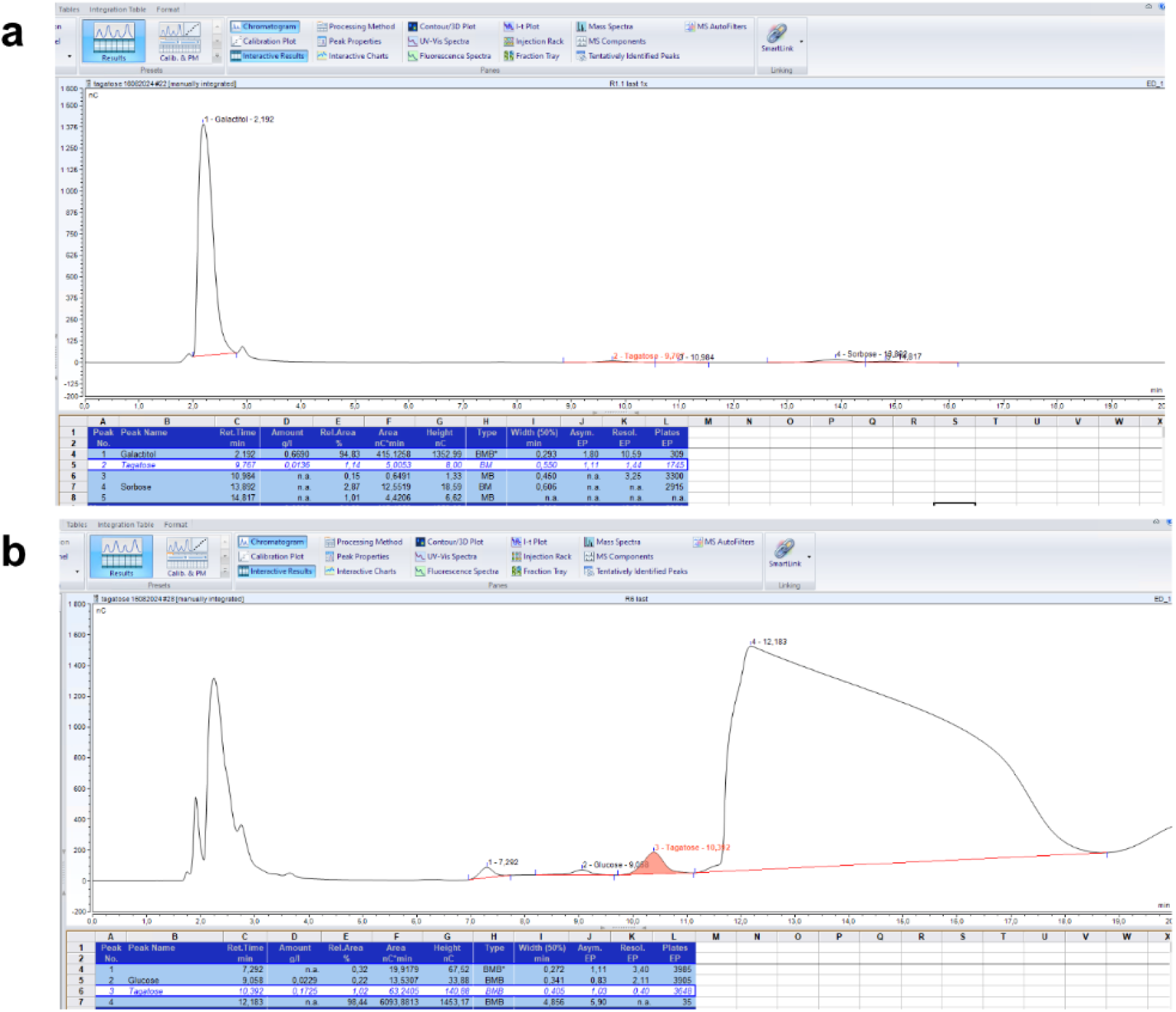
**a** Chromatogram from ion-chromatography analysis of samples from bioreactor run with the cultivation of S. intermedia wild-type strain in minimal media containing 2% galactose as carbon source and 5% ammonium sulphate (High Nitrogen condition) in the media. Culture was maintained at pH 5.5 and with 21% oxygen in the inlet air. Galactose was completely consumed at the endpoint and tagatose and L-sorbose were detected in addition to galactitol in the media. Compounds were detected against internal standards for the respective compounds. **b**: Detection of tagatose in galΔ mutant cultivated in CWP in 1L bioreactor.

**Figure S7.**
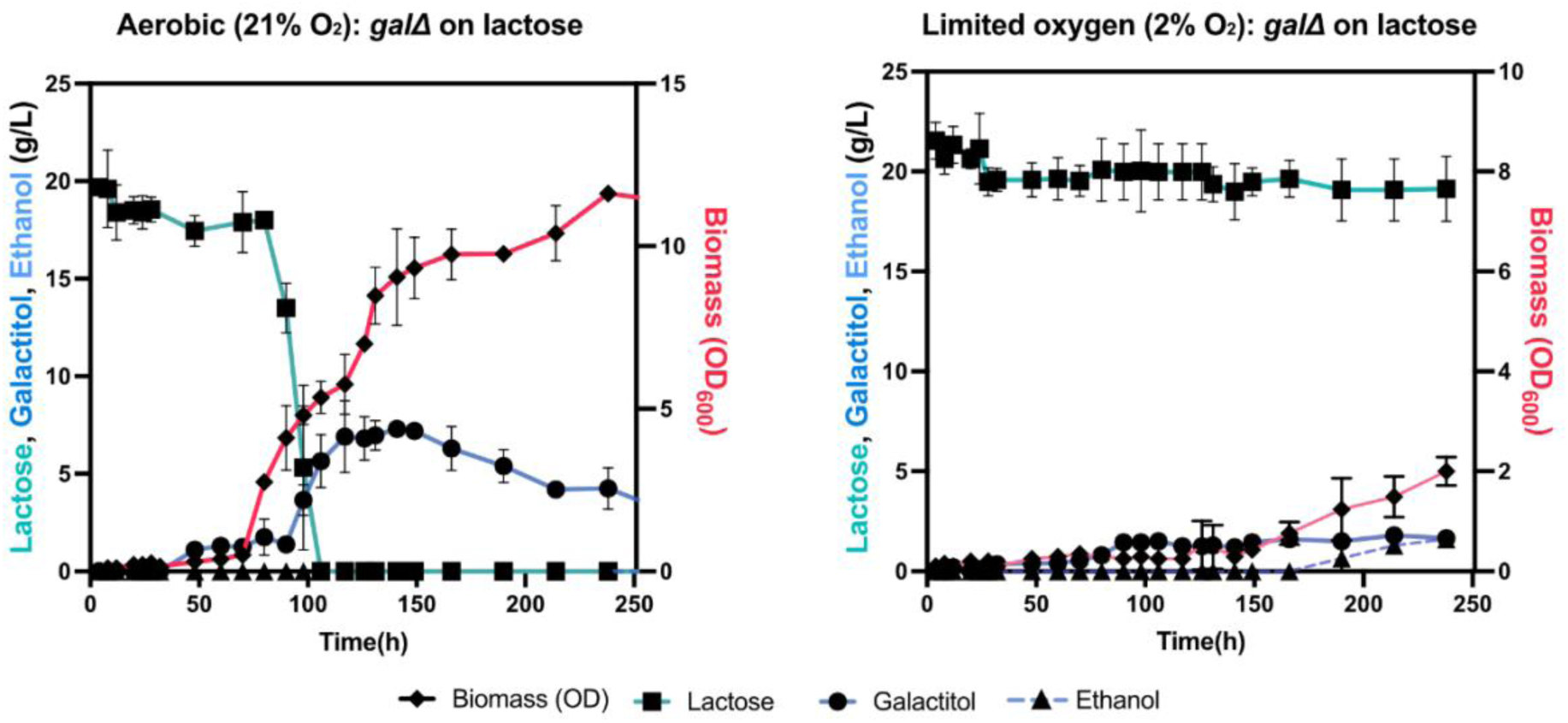
Growth and metabolite production profiles from batch culture in 1L bioreactor for aerobic (21% oxygen in inlet air) vs low-oxygen (2% oxygen in inlet air): Cultivation of gal strain minimal media with 2% lactose as starting substrate. Culture conditions were maintained as mentioned in the materials and methods section except for the percentage of oxygen supplied in the inlet air.

## References

1. Linder T: Making the case for edible microorganisms as an integral part of a more sustainable and resilient food production system. Food Security 2019, 11:265–278.

2. Choi KR, Jung SY, Lee SY: From sustainable feedstocks to microbial foods. Nature Microbiology 2024, 9:1167–1175.

3. Graham AE, Ledesma-Amaro R: The microbial food revolution. Nature Communications 2023, 14:2231.

4. Ledesma-Amaro R, Nicaud JM: Yarrowia lipolytica as a biotechnological chassis to produce usual and unusual fatty acids. vol. 612016.

5. Fraser RZ, Shitut M, Agrawal P, Mendes O, Klapholz S: Safety evaluation of soy leghemoglobin protein preparation derived from *Pichia pastoris*, intended for use as a flavor catalyst in plant-based meat. International journal of toxicology 2018, 37:241–262.

6. Sohn SB, Graf AB, Kim TY, Gasser B, Maurer M, Ferrer P, Mattanovich D, Lee SY: Genome-scale metabolic model of methylotrophic yeast *Pichia pastoris* and its use for in silico analysis of heterologous protein production. Biotechnology Journal 2010.

7. Van Ooyen AJJ, Dekker P, Huang M, Olsthoorn MMA, Jacobs DI, Colussi PA, Taron CH: Heterologous protein production in the yeast *Kluyveromyces lactis*. FEMS Yeast Research 2006, 6:381–392.

8. Liu F, Hu ZD, Zhao XM, Zhao WN, Feng ZX, Yurkov A, Alwasel S, Boekhout T, Bensch K, Hui FL, et al: Phylogenomic analysis of the *Candida auris-Candida haemuli* clade and related taxa in the Metschnikowiaceae, and proposal of thirteen new genera, fifty-five new combinations and nine new species. Persoonia 2024, 52:22–43.

9. Shen XX, Opulente DA, Kominek J, Zhou X, Steenwyk JL, Buh KV, Haase MAB, Wisecaver JH, Wang M, Doering DT, et al: Tempo and Mode of Genome Evolution in the Budding Yeast Subphylum. Cell 2018, 175:1533–1545 e1520.

10. Young EM, Comer AD, Huang H, Alper HS: A molecular transporter engineering approach to improving xylose catabolism in *Saccharomyces cerevisiae*. Metab Eng 2012, 14:401–411.

11. Wu J, Hu J, Zhao S, He M, Hu G, Ge X, Peng N: Single-cell Protein and Xylitol Production by a Novel Yeast Strain *Candida intermedia* FL023 from Lignocellulosic Hydrolysates and Xylose. Appl Biochem Biotechnol 2018, 185:163–178.

12. Nidetzky B, Bruggler K, Kratzer R, Mayr P: Multiple forms of xylose reductase in *Candida intermedia*: comparison of their functional properties using quantitative structure-activity relationships, steady-state kinetic analysis, and pH studies. J Agric Food Chem 2003, 51:7930–7935.

13. Gardonyi M, Osterberg M, Rodrigues C, Spencer-Martins I, Hahn-Hagerdal B: High capacity xylose transport in *Candida intermedia* PYCC 4715. FEMS Yeast Res 2003, 3:45–52.

14. Fonseca C, Olofsson K, Ferreira C, Runquist D, Fonseca LL, Hahn-Hagerdal B, Liden G: The glucose/xylose facilitator Gxf1 from *Candida intermedia* expressed in a xylose-fermenting industrial strain of Saccharomyces cerevisiae increases xylose uptake in SSCF of wheat straw. Enzyme Microb Technol 2011, 48:518–525.

15. Geijer C, Faria-Oliveira F, Moreno AD, Stenberg S, Mazurkewich S, Olsson L: Genomic and transcriptomic analysis of *Candida intermedia* reveals the genetic determinants for its xylose-converting capacity. Biotechnol Biofuels 2020, 13:48.

16. Moreno AD, Carbone A, Pavone R, Olsson L, Geijer C: Evolutionary engineered *Candida intermedia* exhibits improved xylose utilization and robustness to lignocellulose-derived inhibitors and ethanol. Appl Microbiol Biotechnol 2019, 103:1405–1416.

17. Peri KVR, Faria-Oliveira F, Larsson A, Plovie A, Papon N, Geijer C: Split-marker-mediated genome editing improves homologous recombination frequency in the CTG clade yeast *Candida intermedia*. FEMS Yeast Res 2023, 23.

18. Peri KVR, Yuan L, Oliveira FF, Persson K, Alalam HD, Olsson L, Larsbrink J, Kerkhoven EJ, Geijer C: A unique metabolic gene cluster regulates lactose and galactose metabolism in the yeast *Candida intermedia*. Applied and Environmental Microbiology, 0:e01135–01124.

19. Zhai B, Steinø A, Bacha J, Brown D, Daugaard M: Dianhydrogalactitol induces replication-dependent DNA damage in tumor cells preferentially resolved by homologous recombination. Cell Death & Disease 2018, 9:1016.

20. Jiménez-Alcázar M, Curiel-García Á, Nogales P, Perales-Patón J, Schuhmacher AJ, Galán-Ganga M, Zhu L, Lowe SW, Al-Shahrour F, Squatrito M: Dianhydrogalactitol Overcomes Multiple Temozolomide Resistance Mechanisms in Glioblastoma. Mol Cancer Ther 2021, 20:1029–1038.

21. Liu JJ, Zhang GC, Kwak S, Oh EJ, Yun EJ, Chomvong K, Cate JHD, Jin YS: Overcoming the thermodynamic equilibrium of an isomerization reaction through oxidoreductive reactions for biotransformation. Nat Commun 2019, 10:1356.

22. Alper H, Stephanopoulos G: Engineering for biofuels: exploiting innate microbial capacity or importing biosynthetic potential? Nature Reviews Microbiology 2009, 7:715–723.

23. Han T, Nazarbekov A, Zou X, Lee SY: Recent advances in systems metabolic engineering. Current Opinion in Biotechnology 2023, 84:103004.

24. Herrgård MJ, Swainston N, Dobson P, Dunn WB, Arga KY, Arvas M, Büthgen N, Borger S, Costenoble R, Heinemann M, et al: A consensus yeast metabolic network reconstruction obtained from a community approach to systems biology. vol. 26. pp. 1155–11602008: 1155-1160.

25. Nielsen J: Systems Biology of Metabolism. Annual Review of Biochemistry 2017, 86:245–275.

26. Lewis NE, Nagarajan H, Palsson BO: Constraining the metabolic genotype-phenotype relationship using a phylogeny of in silico methods. Nat Rev Microbiol 2012, 10:291–305.

27. Lewis NE, Hixson KK, Conrad TM, Lerman JA, Charusanti P, Polpitiya AD, Adkins JN, Schramm G, Purvine SO, Lopez-Ferrer D, et al: Omic data from evolved *E. coli* are consistent with computed optimal growth from genome-scale models. Molecular Systems Biology 2010, 6.

28. Edwards JS, Palsson BO: The *Escherichia coli* MG1655 in silico metabolic genotype: its definition, characteristics, and capabilities. Proceedings of the National Academy of Sciences 2000, 97:5528–5533.

29. Famili I, Forster J, Nielsen J, Palsson BØ: *Saccharomyces cerevisiae* phenotypes can be predicted by using constraint-based analysis of a genome-scale reconstructed metabolic network. Proceedings of the National Academy of Sciences of the United States of America 2003, 100:13134–13139.

30. Somerville V, Grigaitis P, Battjes J, Moro F, Teusink B: Use and limitations of genome-scale metabolic models in food microbiology. Current Opinion in Food Science 2022, 43:225–231.

31. Agren R, Mardinoglu A, Asplund A, Kampf C, Uhlen M, Nielsen J: Identification of anticancer drugs for hepatocellular carcinoma through personalized genome-scale metabolic modeling. Molecular Systems Biology 2014.

32. Gatto F, Miess H, Schulze A, Nielsen J: Flux balance analysis predicts essential genes in clear cell renal cell carcinoma metabolism. Scientific Reports 2015.

33. Gatto F, Ferreira R, Nielsen J: Pan-cancer analysis of the metabolic reaction network. Metabolic Engineering 2020, 57.

34. Nilsson A, Nielsen J: Genome scale metabolic modeling of cancer. vol. 432017.

35. Lam S, Hartmann N, Benfeitas R, Zhang C, Arif M, Turkez H, Uhlén M, Englert C, Knight R, Mardinoglu A: Systems analysis reveals ageing-related perturbations in retinoids and sex hormones in alzheimer’s and parkinson’s diseases. Biomedicines 2021, 9.

36. Mardinoglu A, Agren R, Kampf C, Asplund A, Uhlen M, Nielsen J: Genome-scale metabolic modelling of hepatocytes reveals serine deficiency in patients with non-alcoholic fatty liver disease. Nature Communications 2014, 5:1–11.

37. Harrison M-C, Ubbelohde EJ, LaBella AL, Opulente DA, Wolters JF, Zhou X, Shen X-X, Groenewald M, Hittinger CT, Rokas A: Machine learning illuminates how diet influences the evolution of yeast galactose metabolism. bioRxiv 2023:2023.2007.2020.549758.

38. Chroumpi T, Martínez-Reyes N, Kun RS, Peng M, Lipzen A, Ng V, Tejomurthula S, Zhang Y, Grigoriev IV, Mäkelä MR, et al: Detailed analysis of the D-galactose catabolic pathways in Aspergillus niger reveals complexity at both metabolic and regulatory level. Fungal Genetics and Biology 2022, 159:103670.

39. Lu H, Li F, Yuan L, Domenzain I, Yu R, Wang H, Li G, Chen Y, Ji B, Kerkhoven EJ, Nielsen J: Yeast metabolic innovations emerged via expanded metabolic network and gene positive selection. Mol Syst Biol 2021, 17:e10427.

40. Lu H, Li F, Sánchez BJ, Zhu Z, Li G, Domenzain I, Marcišauskas S, Anton PM, Lappa D, Lieven C, et al: A consensus *S. cerevisiae* metabolic model Yeast8 and its ecosystem for comprehensively probing cellular metabolism. Nature Communications 2019.

41. Mojzita D, Herold S, Metz B, Seiboth B, Richard P: l-xylo-3-Hexulose Reductase Is the Missing Link in the Oxidoreductive Pathway for d-Galactose Catabolism in Filamentous Fungi*. Journal of Biological Chemistry 2012, 287:26010–26018.

42. Anton M, Almaas E, Benfeitas R, Benito-Vaquerizo S, Blank LM, Dräger A, Hancock JM, Kittikunapong C, König M, Li F, et al: standard-GEM: standardization of open-source genome-scale metabolic models. bioRxiv 2023:2023.2003.2021.512712-512023.512703.512721.512712.

43. Patil KR, Nielsen J: Uncovering transcriptional regulation of metabolism by using metabolic network topology. Proceedings of the National Academy of Sciences 2005, 102:2685–2689.

44. Orth JD, Thiele I, Palsson BØ: What is flux balance analysis? Nature Biotechnology 2010, 28:245–248.

45. Herrmann HA, Dyson BC, Vass L, Johnson GN, Schwartz J-M: Flux sampling is a powerful tool to study metabolism under changing environmental conditions. npj Systems Biology and Applications 2019, 5:32.

46. Neuhauser W, Haltrich D, Kulbe KD, Nidetzky B: NAD(P)H-dependent aldose reductase from the xylose-assimilating yeast *Candida tenuis*. Isolation, characterization and biochemical properties of the enzyme. Biochem J 1997, 326 ( Pt 3):683–692.

47. Watanabe S, Pack SP, Saleh AA, Annaluru N, Kodaki T, Makino K: The positive effect of the decreased NADPH-preferring activity of xylose reductase from *Pichia stipitis* on ethanol production using xylose-fermenting recombinant *Saccharomyces cerevisiae*. Biosci Biotechnol Biochem 2007, 71:1365–1369.

48. Hwang J, Peterson BG, Knupp J, Baldridge RD: The ERAD system is restricted by elevated ceramides. Sci Adv 2023, 9:eadd8579.

49. Verhagen KJA, van Gulik WM, Wahl SA: Dynamics in redox metabolism, from stoichiometry towards kinetics. Current Opinion in Biotechnology 2020, 64:116–123.

50. Domenzain I, Sánchez B, Anton M, Kerkhoven EJ, Millán-Oropeza A, Henry C, Siewers V, Morrissey JP, Sonnenschein N, Nielsen J: Reconstruction of a catalogue of genome-scale metabolic models with enzymatic constraints using GECKO 2.0. Nature Communications 2022, 13:3766.

51. Noor E, Flamholz A, Bar-Even A, Davidi D, Milo R, Liebermeister W: The Protein Cost of Metabolic Fluxes: Prediction from Enzymatic Rate Laws and Cost Minimization. PLOS Computational Biology 2016, 12:e1005167.

52. Robinson SL, Terlouw BR, Smith MD, Pidot SJ, Stinear TP, Medema MH, Wackett LP: Global analysis of adenylate-forming enzymes reveals β-lactone biosynthesis pathway in pathogenic Nocardia. J Biol Chem 2020, 295:14826–14839.

53. Sánchez BJ, Zhang C, Nilsson A, Lahtvee PJ, Kerkhoven EJ, Nielsen J: Improving the phenotype predictions of a yeast genome-scale metabolic model by incorporating enzymatic constraints. Mol Syst Biol 2017, 13:935.

54. Chroumpi T, Peng M, Aguilar-Pontes MV, Muller A, Wang M, Yan J, Lipzen A, Ng V, Grigoriev IV, Makela MR, de Vries RP: Revisiting a ‘simple’ fungal metabolic pathway reveals redundancy, complexity and diversity. Microb Biotechnol 2021, 14:2525–2537.

55. Estrada M, Navarrete C, Møller S, Quirós M, Martínez JL: Open (non-sterile) cultivations of *Debaryomyces hansenii* for recombinant protein production combining industrial side-streams with high salt content. New Biotechnology 2023, 78:105–115.

56. de Jongh WA, Bro C, Ostergaard S, Regenberg B, Olsson L, Nielsen J: The roles of galactitol, galactose-1-phosphate, and phosphoglucomutase in galactose-induced toxicity in *Saccharomyces cerevisiae*. Biotechnol Bioeng 2008, 101:317–326.

57. Verduyn C, Postma E, Scheffers WA, Van Dijken JP: Effect of benzoic acid on metabolic fluxes in yeasts: a continuous-culture study on the regulation of respiration and alcoholic fermentation. Yeast 1992, 8:501–517.

58. Team RC: RA language and environment for statistical computing, R Foundation for Statistical. Computing 2020.

59. Smyth GK: Limma: linear models for microarray data. In Bioinformatics and computational biology solutions using R and Bioconductor. Springer; 2005: 397–420.

60. Engler C, Kandzia R, Marillonnet S: A One Pot, One Step, Precision Cloning Method with High Throughput Capability. PLOS ONE 2008, 3:e3647.

61. Bruder S, Reifenrath M, Thomik T, Boles E, Herzog K: Parallelised online biomass monitoring in shake flasks enables efficient strain and carbon source dependent growth characterisation of *Saccharomyces cerevisiae*. Microb Cell Fact 2016, 15:127.

